# TRMT10A deficiency and tRNA fragmentation disrupt human pancreatic β-cell identity and insulin maturation

**DOI:** 10.64898/2026.07.22.739528

**Authors:** Cristina Moreno-Castro, Maria Nicol Arroyo, Khadija Benabdallah, Ane Olazagoitia Garmendia, Bruno Santacreu, Toshiaki Sawatani, Jonathan Green, Nathalie Pachera, Cristina Cosentino, David Communi, Xavier Bisteau, Virginie Imbault, Soumi Srimani, Jarne Pauwels, Kris Gevaert, Jean-Christophe Jonas, Miriam Cnop, Romano Regazzi, Mariana Igoillo-Esteve

**Affiliations:** ULB Center for Diabetes Research, Université Libre de Bruxelles, Brussels, Belgium; Biobizkaia Health Research Institute, Cruces-Barakaldo, Spain; Department of Biochemistry and Molecular Biology, University of Basque Country (EHU), Leioa, Spain; Department of Fundamental Neurosciences, University of Lausanne, Lausanne, Switzerland; IRIBHM JE Dumont, Université Libre de Bruxelles, Brussels, Belgium; Analytical platform of the Faculty of Pharmacy, Université Libre de Bruxelles Brussels, Belgium; Université catholique de Louvain (UCLouvain), Institute of experimental and clinical research (IREC), Pole of endocrinology, diabetes and nutrition (EDIN), Brussels, Belgium; VIB Center for Medical Biotechnology, VIB, Ghent, Belgium; Department of Biomolecular Medicine, Ghent University, Ghent, Belgium; Division of Endocrinology, ULB Erasmus Hospital, Brussels University Hospital, Université Libre de Bruxelles, Brussels, Belgium; WEL Research Institute, Wavre, Belgium

## Abstract

Mutations in the tRNA-modifying enzyme *TRMT10A* cause a rare monogenic syndrome characterized by early-onset diabetes and neurodevelopmental defects, yet the molecular mechanisms underlying TRMT10A diabetes remain unclear. Using human *TRMT10A*-deficient (knockout and mutant) induced pluripotent stem cells (iPSCs) differentiated into islet-like aggregates and *TRMT10A*-silenced EndoC-βH1 human β-cells, we show that *TRMT10A* deficiency impairs β-cell differentiation, insulin content and glucose-stimulated insulin secretion while inducing widespread transcriptional alterations. These defects are accompanied by oxidative stress, diminished antioxidant capacity, and defective proinsulin processing driven by reduced *PCSK1* expression. Mechanistically, the loss of TRMT10A promotes tRNA fragmentation and the generation of fragments derived from the 5’ end of tRNA^Gln-CTG^ (tDR^Gln-CTG^) that interact with hnRNPM. Our data support the existence of a previously unrecognized hnRNPM–PTBP1 interaction in human β-cells and suggest that this complex may contribute to the regulation of *PCSK1* mRNA stability and/or translation. Furthermore, through its interaction with hnRNPM, tDR^Gln-CTG^ may alter the function of the complex thereby contributing to reduced *PCSK1* expression, defective proinsulin processing, and impaired insulin content. Our findings link tRNA fragmentation, RNA-binding protein networks and insulin maturation, positioning TRMT10A as a critical regulator of β-cell identity and function and uncovering a novel mechanism of β-cell failure in the pathogenesis of diabetes.

## Introduction

Transfer RNAs (tRNAs) are highly posttranscriptionally modified non-coding RNAs best known for their essential role in protein translation. Beyond their canonical role in mRNA decoding, tRNAs and their derivatives regulate diverse cellular processes(Kirchner and Ignatova, 2015; Kuhle et al., 2023; Orellana et al., 2022; Pan, 2018). tRNA modifications, which are introduced by specific tRNA modifying enzymes, are critical for tRNA stability and function(Arroyo et al., 2021; Cosentino et al., 2019; Kirchner and Ignatova, 2015; Orellana et al., 2022; Pan, 2018). Genetic or environmental disruption of tRNAs or their modifying enzymes can lead to abnormal tRNA aminoacylation, modification or fragmentation, ultimately contributing to the pathogenesis of neurological disorders, cancer and diabetes(Arroyo et al., 2021; Bayazit et al., 2022; Brozzi et al., 2024; Cosentino et al., 2019; Jacovetti et al., 2024). tRNA fragmentation refers to the enzymatic cleavage of tRNAs, which can be triggered by multiple factors such as altered modification status of the parental tRNA, cellular stress, or upregulation of particular endonucleases including angiogenin, DICER, ELAC2, and others(Arroyo et al., 2021; Brozzi et al., 2024; Cosentino et al., 2018; Fu et al., 2009; Green et al., 2020; Guzzi et al., 2018; Jacovetti et al., 2024; Kuhle et al., 2023; Kumar et al., 2016; Li et al., 2018; Lu et al., 2022; Maqueda et al., 2023; Mathew et al., 2023; Nientiedt et al., 2016; Peng et al., 2023; Qin et al., 2020; Schultz and Kothe, 2024; Thompson et al., 2008; Wang et al., 2013). This process generates tRNA-derived fragments (tDRs), a novel class of small non-coding RNAs that regulate biological processes by modulating transcription, translation, RNA stability, cell viability, and the localization or function of RNA-binding proteins(Arroyo et al., 2021; Gebetsberger et al., 2017; Kuhle et al., 2023; Kumar et al., 2016; Torres et al., 2019). The function of tDRs is highly context-dependent and influenced by their sequence, posttranscriptional modifications, folding, length, and subcellular localization(Kuhle et al., 2023). Dysregulated tDR levels have been implicated in paternal inheritance of metabolic traits, cancer, neurological disorders, and different forms of diabetes(Bayazit et al., 2022; Brozzi et al., 2024; Cosentino et al., 2018; Jacovetti et al., 2024; Kuhle et al., 2023; Winek and Soreq, 2025).

We and others have demonstrated that biallelic loss-of-function mutations in *TRMT10A,* encoding the tRNA methyltransferase TRMT10A, cause a rare autosomal recessive syndrome characterized by non-autoimmune early-onset diabetes mellitus, microcephaly, intellectual disability, epilepsy and short stature(Brener et al., 2022; Ceraolo et al., 2025; Gillis et al., 2014; Igoillo-Esteve et al., 2013; Lin et al., 2020; Narayanan et al., 2015; Samhani et al., 2024; Siklar et al., 2021; Stern et al., 2021; Yew et al., 2016; Zung et al., 2015). In most *TRMT10A*-deficient individuals, diabetes manifests in adolescence or early adulthood and, in some cases, it is preceded by hyperinsulinemic hypoglycemia(Brener et al., 2022; Ceraolo et al., 2025; Gillis et al., 2014; Siklar et al., 2021; Zung et al., 2015). TRMT10A catalyzes methylation of specific cytosolic tRNAs at guanosine in position 9 (m^1^G_9_)(Cosentino et al., 2018; Howell et al., 2019; Igoillo-Esteve et al., 2013; Vilardo et al., 2020). Using various models of *TRMT10A* deficiency we previously showed that the loss of TRMT10A induces oxidative stress and activates the intrinsic pathway of apoptosis in rat and human pancreatic β-cells, leading to increased basal and stress-induced β-cell death(Cosentino et al., 2018; Igoillo-Esteve et al., 2013). Notably, we demonstrated that in lymphoblasts and induced pluripotent stem cell (iPSC)-derived β-cells from a *TRMT10A*-deficient individual, the TRMT10A substrate tRNA^Gln-CTG^ undergoes fragmentation, and the resulting tDR^-Gln-CTG^ contributes to β-cell death(Cosentino et al., 2018). Despite these insights, the pathogenic mechanisms of TRMT10A diabetes remain largely unknown.

In this study, we used *TRMT10A* knockout (KO) and mutant human β-like cells to investigate the impact of *TRMT10A* deficiency and tDR^-Gln-CTG^ formation on β-cell development and function and charted the underlying gene and protein expression patterns.

## Results

### TRMT10A deficiency alters β-cell development and fate

The *TRMT10A* gene was knocked out in the previously validated control human iPSC line 1023A(Lytrivi et al., 2021) using CRISPR/Cas9 technology(Virgilio et al., 2025). Two guide (g)RNAs targeting *TRMT10A* exon 3 and exon 4 (Supplementary Table 1) were designed to induce a 724-nucleotide deletion, causing a frameshift and a premature stop codon. However, the guides bound asynchronously, resulting in independent gene re-arrangements. gRNA#1 caused the loss of 2 amino acids in exon 3 without altering the reading frame, whereas gRNA#2 introduced a premature stop codon in exon 4 in both alleles. Absence of TRMT10A protein was confirmed in the KO iPSC line by Western blot (Fig. 1A). The c.379G>A (p.Arg127Stop) *TRMT10A* mutation present in the previously validated iPSC line (HEL122.2, henceforth called *TRMT10A* mutant) from a *TRMT10A*-deficient individual(Cosentino et al., 2018) was corrected using CRISPR-Cas9. This was achieved with gRNA#2 and a donor template designed to repair the mutation and restore the reading frame, generating two isogenic corrected clones. The restoration of TRMT10A expression was confirmed by Western blot (Fig. 1B). *TRMT10A* corrected and *TRMT10A* KO iPSCs were devoid of chromosomal abnormalities (Supplementary Fig. 1A and B) and off target editing (Supplementary Tables 2 and 3), expressed the pluripotency markers OCT4, SSEA4, Nanog and TRA1-60 (Supplementary Fig. 1C), and were able to differentiate into all 3 germ layers (Supplementary Fig. 1D), validating their pluripotency. The *TRMT10A*-deficient iPSC lines and isogenic controls were then differentiated into islet-like aggregates using a previously validated 7-stage protocol(Virgilio et al., 2025) (Fig. 1C). At stage 1 (St1), SRY-box transcription factor 17 (SOX17) staining was comparable in *TRMT10A*-deficient and control lines indicating successful definitive endoderm commitment (Supplementary Fig. 2A). At stage 4 (St4), pancreatic and duodenal homeobox 1 (PDX1) staining remained similar in isogenic control and *TRMT10A*-deficient cells but NK6 homeobox 1 (NKX6.1) was noticeably weaker in *TRMT10A* KO and especially in *TRMT10A*-mutant cells (Fig. 1d and Supplementary Fig. 2B). In line with this, *NKX6.1* mRNA expression was significantly reduced in *TRMT10A* KO St7 islet-like aggregates while no differences in the expression of the additional differentiation markers were observed (Supplementary Fig. 3). St7 *TRMT10A*-deficient 3D islet-like aggregates appeared smaller than control islets in bright field (Fig. 1E and Supplementary Fig. 2C). By immunofluorescence analysis of St7 cells, the proportion of β-like cells in *TRMT10A* KO islet-like aggregates was comparable to that of control aggregates (Fig. 1F), but *TRMT10A*-mutant aggregates displayed a significantly lower β-like cell proportion than their isogenic corrected counterparts (Fig. 1G), consistent with the marked reduction in NKX6.1 expression observed at earlier differentiation stages The proportion of α-, δ- and polyhormonal cells was unaffected (Fig. 1E-G).

**Fig. 1.**
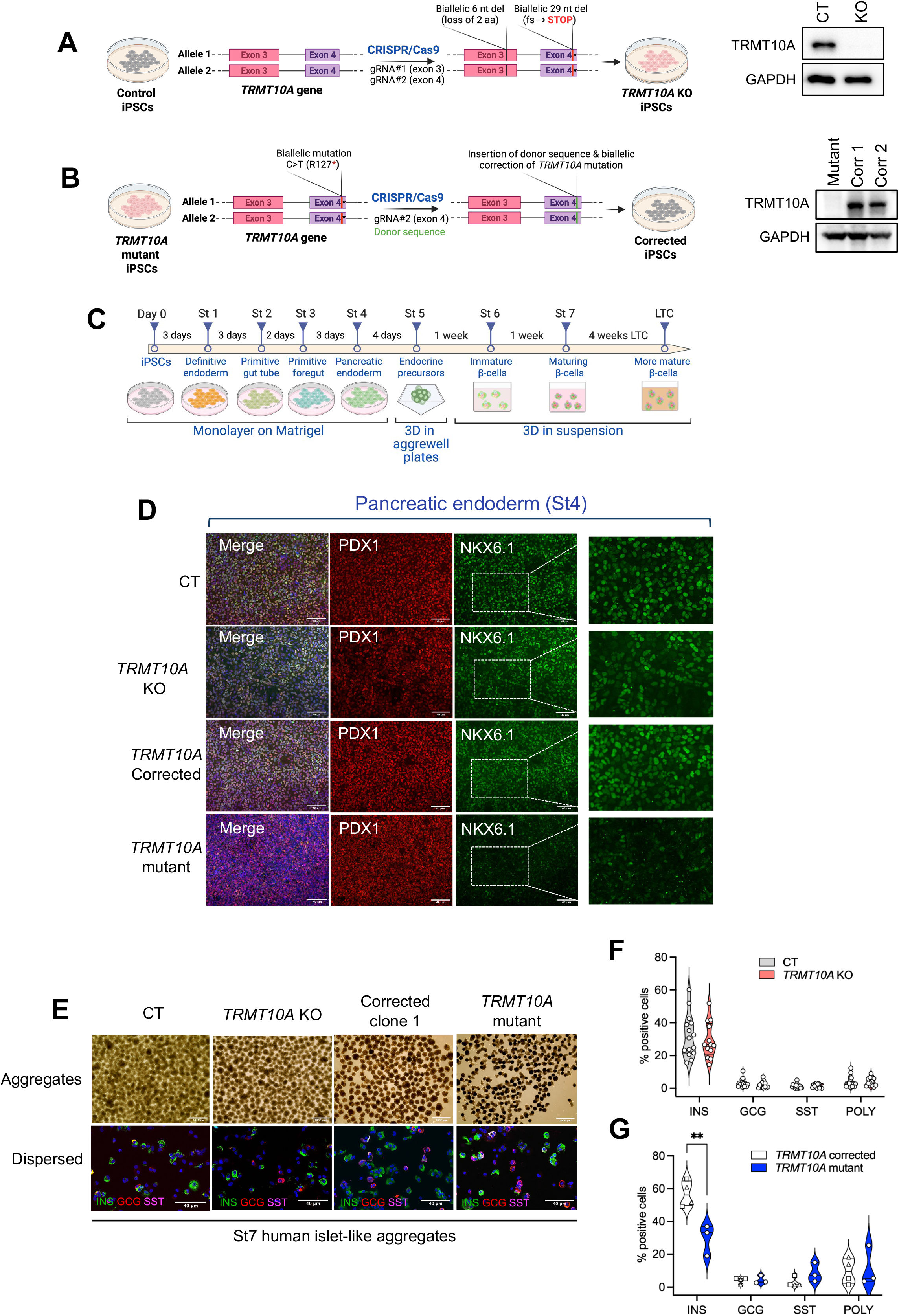
CRISPR-Cas9-mediated TRMT10A gene editing in human iPSCs and their differentiation into human islet-like aggregates. A and. **B**, Schematic representation of the CRISPR-Cas9 approach used to either **A**, knockout (KO) the TRMT10A gene in a control iPSC line or **B**, correct the TRMT10A c.379G>A (p.Arg127Stop) mutation in a TRMT10A mutant iPSC line. **A**, gRNA#1 and gRNA#2 were used to generate the TRMT10A KO. gRNA#1 caused the biallelic loss of 2 aminoacids (aa) in exon 3 without change in the reading frame, and gRNA#2 caused the appearance of a biallelic premature stop codon in exon 4. **B**, gRNA#2 targeted exon 4 at the mutant site, inducing a biallelic cut and enabling insertion of a donor template designed to repair the mutation and restore the open reading frame. The absence of TRMT10A protein in the KO line and the rescue of TRMT10A expression in the two corrected clones was confirmed by Western blot (Mutant: TRMT10A mutant line, Corr 1: corrected clone 1, Corr 2: corrected clone 2). **C**, Schematic representation of the iPSC differentiation protocol. The different stages (St) of the differentiation are highlighted. **D**, Representative images of pancreatic endoderm showing NKX6.1 and PDX1 positivity in isogenic control (CT), TRMT10A KO, TRMT10A mutant and TRMT10A-corrected lines. Scale bar, 40 μm. The pictures on the right are magnifications of the NKX6.1 staining in the highlighted rectangles. **E**, Bright field representative images of 3D aggregates at the end of the differentiation (St7). Scale bar, 1000 μm. Immunofluorescence images of dispersed aggregates immunostained with anti-insulin (INS, green), anti-glucagon (GCG, red), and anti-somatostatin (SST, magenta) antibodies. Scale bar, 40 μm. **F-G**, Proportion of β-cells (INS positive), α-cells (GCG positive), δ-cells (SST positive) and polyhormonal cells (POLY) in St7 islet-like aggregates derived from CT and TRMT10A KO **(F),** or TRMT10A corrected and TRMT10A mutant iPSCs **(G)**. Individual data points represent independent differentiations. In panel **G**, squares correspond to TRMT10A corrected clone 1, and triangles to TRMT10A corrected clone 2. **p<0.01 by Two-way Anova with Šídák’s correction for multiple comparisons. Panels A to C were created with BioRender (https://BioRender.com).

### Insulin processing, content, and secretion are impaired in TRMT10A-deficient β-like cells

To assess the impact of *TRMT10A* deficiency on β-cell function, we measured insulin secretion in *TRMT10A* KO, *TRMT10A* mutant and isogenic control islet-like aggregates. Insulin secretion and content were normalized to the β-cell proportion to account for different β-cell yields in iPSC differentiations. *TRMT10A* KO β-like cells at St7 displayed blunted insulin secretion in response to 16.8 mM glucose, and diminished secretion in response to 16.8 mM glucose plus the sulfonylurea gliclazide, that promotes insulin release by closing K_ATP_ channels, leading to membrane depolarization (Fig. 2A-B). Insulin secretion stimulated by 16.8 mM glucose plus KCl was unaffected. The K_ATP_ channel opener diazoxide suppressed insulin secretion in *TRMT10A* competent and KO β-like cells, confirming that K_ATP_ channels are functional (Fig. 2A-B). Insulin content was significantly reduced in *TRMT10A* KO St7 islet-like aggregates (Fig. 2E). A similar trend of impaired insulin secretion and content was observed in *TRMT10A*-mutant β-like cells relative to their isogenic corrected controls (Fig. 2C, D and F). However, the small number of biological replicates for the mutant cells (n = 2) prevented formal statistical evaluation.

**Fig. 2.**
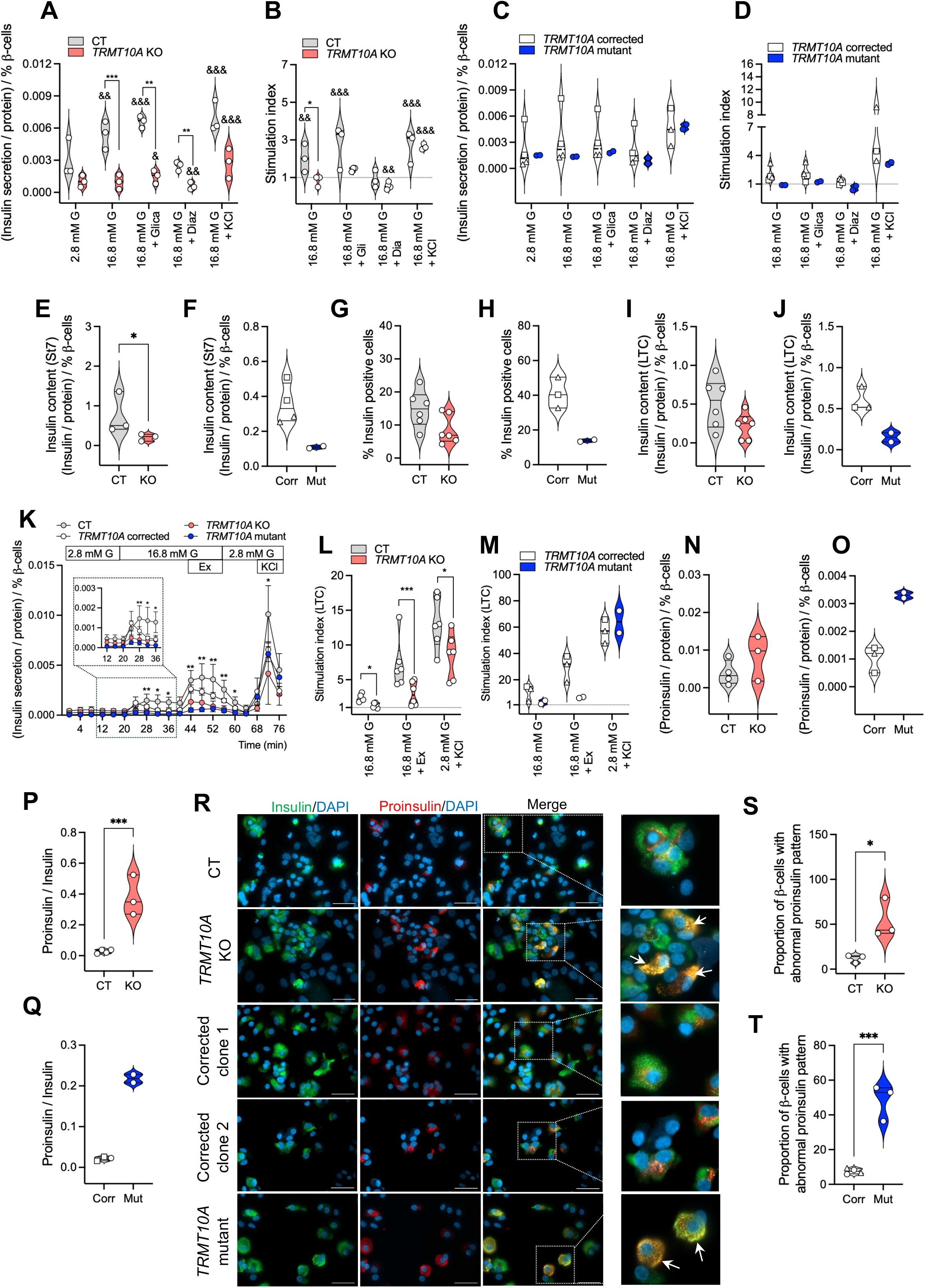
TRMT10A deficiency impairs insulin secretion and content in iPSC-derived β-like cells. **A-D**, Non-sequential static insulin secretion in St7 islet-like aggregates derived from isogenic control (CT) and *TRMT10A* KO (KO) iPSCs **(A-B)**, or *TRMT10A* corrected (Corr) and *TRMT10A* mutant (Mut) iPSCs **(C-D)**. **A** and **C**, Basal, 2.8 mM Glucose (2.8 mM G) and stimulated insulin secretion in response to 16.8 mM glucose alone (16.8 mM G) or combined with 25 μM gliclazide (Gli), 250 μM diazoxide (Dia), and 30 mM KCl (KCl). Data was normalized to total protein and to the β-cell proportion in the aggregates. **B** and **D**, Stimulation index calculated as the ratio between insulin secretion in the different conditions and 2.8 mM glucose. **E-F**, Mean insulin content of St7 islet-like aggregates normalized to their total protein content and β-cell proportion. **G-M**, Data derived from long term cultured (LTC) iPSC-derived islet-like aggregates used for dynamic insulin secretion. **G-H**, Percent of insulin positive cells, assessed by immunofluorescence. **I-J**, Insulin content assessed by ELISA. **K**, Dynamic insulin secretion (n=6 CT, n=6 *TRMT10A* KO, n=3 *TRMT10A* corrected, and n=2 *TRMT10A* mutant differentiations) at 2.8 mM glucose, or 16.8 mM glucose alone or combined with 50 ng/ml Exendin-4 (Ex), or 50 mM KCl. The response to glucose is magnified in the insert. **L-M,** Stimulation index calculated from panel **K**. **N-O,** Proinsulin content in St7 aggregates assessed by ELISA. **P-Q,** Proinsulin to insulin ratio in St7 aggregates. **R**, Immunofluorescence of dispersed St7 aggregates stained for proinsulin (red) and insulin (green). Nuclei were counterstained with DAPI (blue). Representative images from n=6-7 independent differentiations. Scale bars, 40 µm. Higher-magnification views of the highlighted regions are displayed on the right. Arrows indicate cells exhibiting mislocalized proinsulin. **S-T,** Quantification of the pictures. The proportion of cells showing abnormal proinsulin localization with respect to the total β-cell number is shown. Symbols represent independent differentiations. Squares correspond to *TRMT10A* corrected clone 1, and triangles to *TRMT10A* corrected clone 2. * p<0.05, ** p<0.01, *** p<0.001, TRMT10A KO vs CT, & p<0.05, && p<0.01, 2.8 mM glucose vs other conditions by two-way ANOVA followed by Šidák’s correction for multiple comparisons **A, B, K**, and **L**) or by unpaired t-test (**C, P, S** and **T**).

Mitochondrial function, assessed by Seahorse analysis of dispersed St7 aggregates, showed that FCCP-mediated oxygen consumption rate (OCR) was significantly reduced in *TRMT10A* KO cells compared with control (Supplementary Fig. 4A), as was spare respiratory capacity, proton leak and non-mitochondrial respiration, while ATP production was increased (Supplementary Fig. 4B-F). No difference in glucose-induced respiration and maximal respiratory capacity was observed (Supplementary Fig. 4G and H). The diminished spare respiratory capacity of *TRMT10A*-deficient cells points to a decreased ability to respond to elevated energetic demands and stressful conditions making the cells more susceptible to cell death(Armstrong et al., 2018; Sriskanthadevan et al., 2015), while the reduced proton leak may generate production of reactive oxygen species (ROS) and oxidative stress(Cadenas, 2018), a feature that we previously reported in *TRMT10A*-silenced β-cells(Cosentino et al., 2018). Consistent with these findings, NAD(P)H autofluorescence, which was elevated by glucose and reduced by FCCP in St7 control and *TRMT10A* KO islet aggregates, was 43% lower in the latter indicating persistent oxidative stress (Supplementary Fig. 4I). To generate more functionally mature β-like cells, we applied a long-term culture (LTC) differentiation method(Balboa et al., 2022; Virgilio et al., 2025). LTC *TRMT10A* deficient aggregates tended to exhibit a reduced proportion of β-like cells and lower insulin content compared with their isogenic counterparts (Fig. 2G-J). Dynamic insulin secretion assays revealed blunted secretory responses to 16.8 mM glucose, either alone or combined with the GLP-1 receptor (GLP1R) agonist exendin-4 in both, *TRMT10A* KO and *TRMT10A* mutant LTC aggregates (Fig. 2K) with around 40-80% reduced stimulation indexes (Fig. 2L-M).

To examine whether insulin processing is affected by *TRMT10A* deficiency, we quantified proinsulin levels by ELISA. While the proinsulin content was similar between *TRMT10A* KO and isogenic control St7 aggregates (Fig. 2N), the proinsulin-to-insulin ratio was significantly higher under *TRMT10A* deficiency (Fig. 2P). In line with this, the *TRMT10A* mutant aggregates showed a trend for increased proinsulin content and insulin to proinsulin ratio respect to the corrected aggregates (Fig. 2O and Q) pointing to a proinsulin processing defect. Immunostaining of dispersed St7 aggregates revealed that, in control β-cells, proinsulin was primarily localized to the perinuclear region and rarely colocalized with mature insulin (Fig. 2R), while *TRMT10A*-deficient β-cells displayed widespread cytosolic proinsulin distribution, which predominantly colocalized with insulin in 48-54% of β-like cells (Fig. 2R-T). This abnormal pattern, reflecting impaired proinsulin processing, has also been reported in β-cells from individuals with type 1 diabetes diagnosed before the age of seven years(Sims et al., 2019a; Sims et al., 2019b).

Collectively, these findings indicate that *TRMT10A*-deficient β-cells exhibit profound dysfunction, characterized by defective proinsulin processing, reduced insulin content, and impaired insulin secretion. This is accompanied by reduced NAPDH levels and a mitochondrial function profile consistent with heightened oxidative stress.

### TRMT10A deficiency leads to widespread dysregulation of gene and protein expression

To investigate the underlying molecular mechanisms of β-cell dysfunction in *TRMT10A* deficiency, we performed bulk RNA sequencing of St7 aggregates (n=8 control and n=8 *TRMT10A* KO differentiations) and parallel proteomic analyses (n=7). The proportion of β, α, δ and polyhormonal cells in iPSC-derived islets used for these omics is shown in Fig. 3A. The RNA sequencing revealed 1937 differentially expressed (DE) coding and non-coding genes, 917 down- and 1020 upregulated in *TRMT10A* KO vs control (Fig. 3B). From the 6161 proteins detected by proteomics, 420 were DE (237 down- and 183 upregulated, Fig. 3C). Database for Annotation, Visualization and Integrated Discovery (DAVID) functional gene ontology (GO) analysis of DE genes and proteins revealed multiple dysregulated biological processes: extracellular matrix organization, cell-cell adhesion, cell-substrate adhesion and cholesterol homeostasis pathways were upregulated in *TRMT10A*-deficient cells; signal transduction, synaptic vesicle transport, exocytosis, Golgi lumen acidification, insulin secretion, glucose homeostasis, nervous system development and antiviral defense were downregulated (Fig. 3D).

**Fig. 3.**
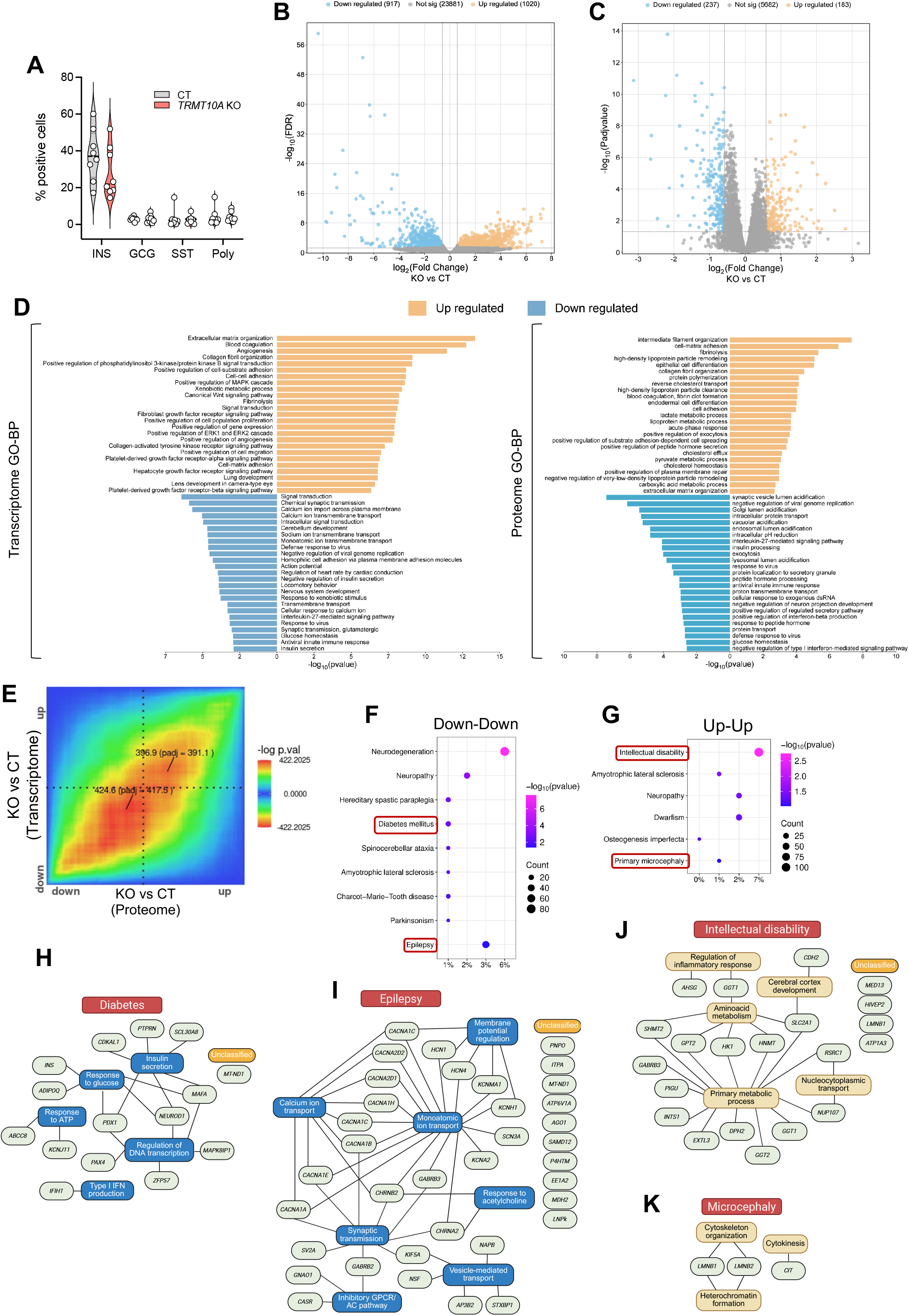
TRMT10A deficiency alters gene and protein expression in iPSC-derived islet-like aggregates. **A**, Proportion of insulin (INS), glucagon (GCG), somatostatin (SST), and polyhormonal (Poly) cells in isogenic control (CT) and *TRMT10A* KO St7 iPSC-derived islet-like aggregates used for bulk RNA sequencing (n = 8) and proteomics (n = 7). **B** and **C,** Volcano plots showing differential expressed (DE) genes **B**, and proteins **C**. Genes and proteins with an absolute log2 fold change (|log2FC|) ≥ 0.585 and false discovery rate (FDR) or adjusted p-value < 0.05 were considered DE. Orange indicates upregulated; blue indicates downregulated. **D**, DAVID functional gene ontology (GO) analysis of DE genes and proteins. The top 25 upregulated and downregulated biological processes (BP) in *TRMT10A* KO cells relative to CT are shown. **E**, Rank–rank hypergeometric overlap map generated using RedRibbon. The heatmap indicates the significance (−log10 p-value) of overlap between proteomic and transcriptomic datasets of *TRMT10A* KO St7 islet-like aggregates. Quadrants represent genes and proteins changing in the same direction (up–up; down–down) or in opposite directions (up–down; down–up). **F-G**, List of diseases (UniProt keyword disease annotations) associated with the down–down and up-up gene sets from RedRibbon. **H-I**, Significantly downregulated genes associated with diabetes and epilepsy and their corresponding BP; **J-K**, Significantly upregulated genes associated with intellectual disability and microcephaly and their corresponding BP. Panels **H** to **K** were created with BioRender (https://BioRender.com).

To compare transcriptomes and proteomes by rank–rank hypergeometric overlap we used RedRibbon(Piron et al., 2024) (Fig. 3E). The heatmap illustrates the significance of overlap between genes that are upregulated in both datasets (top right quadrant, “up-up”), downregulated in both (bottom left quadrant, “down-down”), or oppositely regulated (top left and bottom right quadrants). RedRibbon revealed substantial concordance between the two datasets with 1541 genes commonly up- and 1463 commonly downregulated, and no significant anticorrelation. By GO analysis the “up-up” genes were enriched in pathways related to mRNA splicing regulation, mRNA and rRNA processing, and DNA repair, while the “down-down” genes were involved in protein transport, endoplasmic reticulum (ER) to Golgi vesicle-mediated transport, Golgi organization, exocytosis and endocytosis (Supplementary Fig. 5). By UniProt keyword disease annotation the “down-down” genes were linked to neurodegenerative processes, diabetes and epilepsy, the latter two being clinical features of *TRMT10A*-deficient individuals(Brener et al., 2022; Cosentino et al., 2018; Gillis et al., 2014; Howell et al., 2019; Igoillo-Esteve et al., 2013; Lin et al., 2020; Narayanan et al., 2015; Samhani et al., 2024; Siklar et al., 2021; Stern et al., 2021; Tresky et al., 2024; Yew et al., 2016; Zung et al., 2015) (Fig. 3F). The “up-up” genes were associated with neurological disorders including intellectual disability and primary microcephaly, also features of *TRMT10A*-deficient patients(Brener et al., 2022; Ceraolo et al., 2025; Gillis et al., 2014; Igoillo-Esteve et al., 2013; Lin et al., 2020; Narayanan et al., 2015; Samhani et al., 2024; Siklar et al., 2021; Stern et al., 2021; Yew et al., 2016; Zung et al., 2015) (Fig. 3G). GO analyses of the significantly dysregulated genes of these four features are shown in Fig. 3H-K. Diabetes-related genes were involved in DNA transcription, insulin secretion, β-cell metabolism, glucose sensing and type I interferon production(Deshmukh et al., 2021; Zhu et al., 2017) (Fig. 3H), while epilepsy-related genes encoded voltage-gated calcium channel subunits, ion channels, and proteins related with vesicle transport, membrane potential regulation and synaptic transmission, all key components of neuronal signaling, excitability and neurotransmitter release(Simms and Zamponi, 2014; Trus and Atlas, 2024) (Fig. 3I). Reduced expression of some of these genes was confirmed by RT-qPCR in *TRMT10A* KO St7 islet-like aggregates (Supplementary Fig. 6A). These included *PTPRN,* important for insulin granule biogenesis and turnover(Toledo et al., 2023; Yasui et al., 2024); *MAFA and PAX4*, key regulators of β-cell function(Lau et al., 2023; Wang et al., 2007; Zhu et al., 2017); *ABCC8,* encoding the SUR1 subunit of the K_ATP_ channel(Ashcroft et al., 1984; Rorsman and Trube, 1985); and *IFIH1* which encodes the viral sensor MDA5(del Toro Duany et al., 2015). *CACNA1E* and *NPTX2*, involved in synaptic plasticity, learning and memory were also reduced(Kessi et al., 2021; van der Ende et al., 2020; Xiao et al., 2017; Yang et al., 2024; Zhou et al., 2023) (Supplementary Fig. 6A). *TRMT10A*-mutant islet-like aggregates exhibited a similar trend of reduced expression compared to *TRMT10A* correct cells, although the limited number of biological replicates precluded adequate statistical evaluation (Supplementary Fig. 6, white and blue graphic). Several genes associated with proton transmembrane transport were downregulated (Supplementary Fig. 6B), including UCP2 that inhibits ROS production by catalyzing proton leak across the mitochondrial inner membrane, in keeping with the Seahorse data (Supplementary Fig. 4A)(Zhao et al., 2019).

The upregulated genes associated with intellectual disability were primarily involved in metabolic processes, nucleocytoplasmic transport, cerebral cortex development and inflammatory responses (Fig. 3J), while the microcephaly genes play a role in cytoskeleton organization, heterochromatin formation and cytokinesis (Fig. 3K). Because the iPSC-islet-like aggregates contain a mixed cell population, we compared by RedRibbon the *TRMT10A* KO St7 aggregates proteome with the proteome of clonal human EndoC-βH1 β-cells transfected with two previously validated human *TRMT10A* siRNAs*(Cosentino et al., 2018)*. This revealed 1064 commonly down- and 172 commonly upregulated proteins (Supplementary Fig. 7). GO analysis of the “up-up” and “down-down” proteins confirmed biological processes identified in *TRMT10A* KO St7 islet-like aggregates (Supplementary Fig. 5). These results demonstrate that the transcriptomic and proteomic analyses of *TRMT10A*-null iPSC-derived islet-like aggregates successfully captured biologically relevant β-cell processes.

Overall, these results point to an important transcriptional dysregulation in *TRMT10A*-deficient islets potentially compromising β-cell and neuronal function. They also highlight the pertinence of CRISPR/Cas9-edited iPSC-islet-like models to study TRMT10A disease pathogenesis.

### PCSK1 expression is reduced in TRMT10A-deficient β-like cells

Given the proinsulin processing defects of iPSC-derived *TRMT10A-*deficient β-cells (Fig. 2P-T), we examined the expression of genes directly or indirectly involved in proinsulin processing in the transcriptomic and proteomic datasets. Proinsulin processing is a multistep process occurring in the trans-Golgi network and secretory granules, requiring proper proinsulin folding and tightly regulated pH and calcium conditions(Zavarzadeh et al., 2025). Prohormone convertases, the key enzymes involved in this process, are calcium-dependent serine endoproteases that acquire full catalytic activity in the acidic environment of secretory granules(Rouille et al., 1995). Notably, *PCSK1 (*coding for prohormone convertase 1/3, PC1/3), the primary enzyme for proinsulin cleavage in human β-cells(Ramzy et al., 2020; Stijnen et al., 2016; Torchio et al., 2025), was reduced by more than 4-fold at mRNA and protein level in *TRMT10A-*deficient cells (Fig. 4A). RT-qPCR and Western blot confirmed the decrease in *PCSK1* mRNA and PC1/3 protein levels in *TRMT10A* KO cells, as well as in *TRMT10A* mutant St7 islet-like aggregates (Fig. 4B-G). *TRMT10A* silencing by RNA interference in clonal EndoC-βH1 β-cells also showed reduced PC1/3 expression (Fig. 4H-J). *PCSK2,* coding for prohormone convertase 2, which plays a minimal or context-dependent role in human proinsulin processing, was not detected in the proteomics but reduced by 2-fold at mRNA level. Carboxypeptidase E (CPE), responsible for final proinsulin trimming(Ramzy et al., 2020; Stijnen et al., 2016; Torchio et al., 2025), and the ER oxidoreductase 1 beta (ERO1B), an ER-resident enzyme involved in proinsulin maturation, were reduced by 1.4 to 2-fold at mRNA and protein level (Fig. 4A). Protein disulfide isomerases (*PDIA3-6, P4HB and ERP44)*, critical for disulfide bond formation and proper proinsulin folding, were unchanged. *ATP6AP1*, one of the accessory subunits of the V-ATPase required for granule acidification, was reduced by 1.6-fold in *TRMT10A*-deficient cells suggesting that impaired acidification may contribute to the insulin maturation defect (Fig. 4A). *ATP2A2* expression, encoding SERCA2, essential for ER calcium homeostasis, was unaffected by the lack in TRMT10A, while *ATP2A3*, encoding SERCA3, was reduced by 2-fold (Fig. 4A). Collectively, *TRMT10A*-deficient cells have gene and protein signatures consistent with defective proinsulin processing with predominant loss of *PCSK1* (PC1/3).

**Fig. 4:**
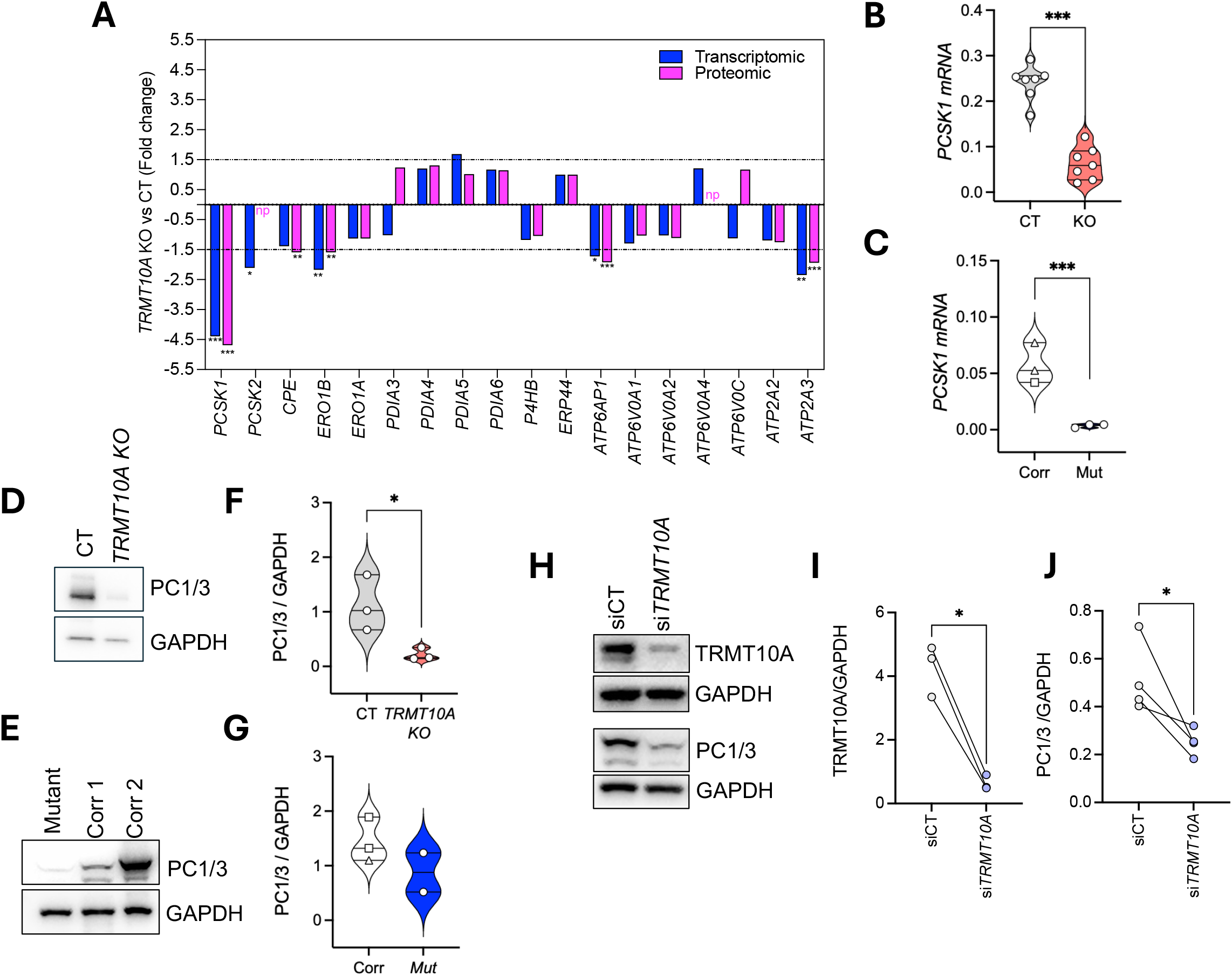
PCSK1 expression is reduced in TRMT10A-deficient cells. **A**, Transcriptomic (blue) and proteomic (pink) profiles of genes directly or indirectly involved in proinsulin processing in pancreatic β-cells. Differential expression is shown as fold change. Statistical significance was defined as FDR or adjusted p-value < 0.05. np=not present in the proteomic dataset. **B** and **C**, *PCSK1* mRNA expression assessed by RT-qPCR in **B**, isogenic control (CT) and *TRMT10A* KO or **C**, *TRMT10A* corrected (Corr) and *TRMT10A* mutant (Mut) St7 islet-like aggregates. mRNA expression was normalized to *ACTB* and expressed as 2^−ΔCt^. **D-G**, Expression of protein convertase 1/3 (PC1/3) in CT and *TRMT10A* KO **(D** and **F)** or *TRMT10A* corrected (Corr) and *TRMT10A* mutant (Mut) St7 islet-like aggregates **(E** and **G)**. **H-J** expression of PC1/3 in *TRMT10A*-silenced EndoC-βH1 cells (si*TRMT10A*). **D, E** and **H**, Representative Western blot images of 2–4 independent experiments. **F, G, I** and **J**, Quantification of the blots. PC1/3 and TRMT10A protein levels were normalized to GAPDH. Data points represent independent differentiations or silencing experiments. Squares correspond to *TRMT10A* corrected clone 1, and triangles corrected clone 2. *p<0.05, *** p<0.001 by unpaired t-test (**B, C** and **F)**, or ratio paired t-test (**I** and **J).**

### tRNA^Gln^ fragmentation in TRMT10A-deficient human β- and non β-cells

Using lymphoblasts and iPSC-derived St7 islet-like aggregates from a *TRMT10A*-deficient individual and non-isogenic controls, we had previously demonstrated that TRMT10A deficiency is associated with increased fragmentation of tRNA^Gln-CTG^ (Cosentino et al., 2018). In keeping with these earlier findings, ALKB + AlkB-D135S-demethylated RNA(Cosentino et al., 2018; Zheng et al., 2015) from *TRMT10A-*silenced EndoC-βH1 cells (Fig. 5A) and *TRMT10A* KO St7 islet aggregates showed increased levels of two fragments, namely tDR^-1:18-Gln-CTG-1-M10^ (henceforth called tDR^-1:18-Gln^, Fig. 5B-D) and tDR^-1:29-Gln-CTG-1-M3^ (henceforth called tDR^-1:29-Gln^, Fig. 5E-G). The levels of the parental tRNA tRNA^Gln-CTG^ were unchanged (Fig. 5H-I). As an orthogonal approach, we developed a ligation-based Fluorescent In-Situ Hybridization technique (Li-FISH) to specifically detect a tDR of interest without detecting its parental tRNA. This approach, schematized in Fig. 5J, is not affected by the m^1^G_9_ modification of the tDR since the probe anneals downstream of G_9_. In good concordance with the RT-qPCR results, Li-FISH showed increased tDR^-1:29-Gln^ in *TRMT10A* KO St7 β- and non-β cells (Fig. 5K) and *TRMT10A-*silenced EndoC-βH1 cells (Supplementary Fig. 8). In both models tDR^-1:29-Gln^ appeared enriched in the cytoplasm. In line with the Li-FISH findings, subcellular RNA fractionation followed by RT-qPCR in *TRMT10A* KO St7 aggregates showed mostly cytoplasmic localization of tDR^-1:29-Gln^ and predominant nuclear localization of the shorter tDR^-1:18-Gln^ (Fig. 5L).

**Fig. 5:**
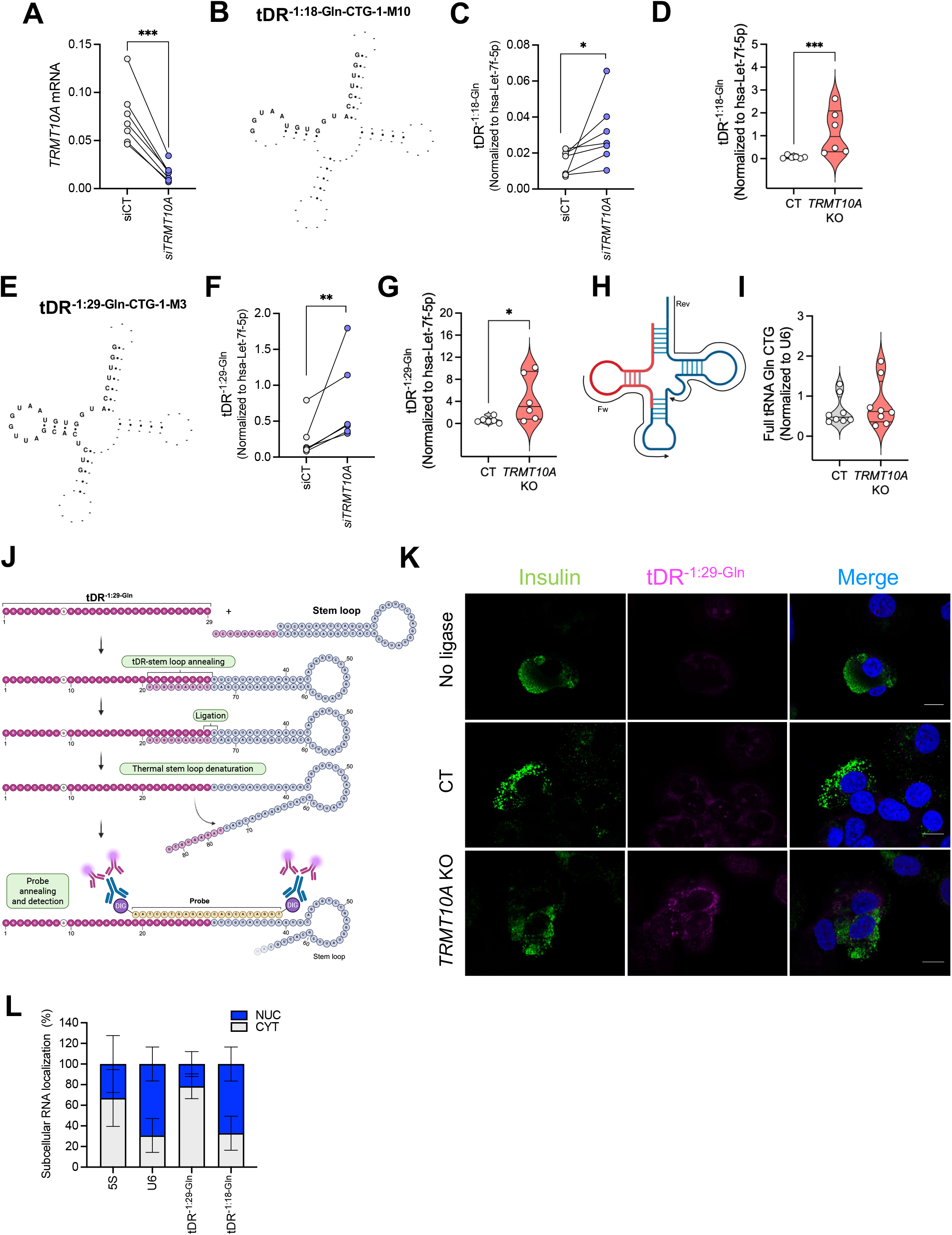
TRMT10A deficiency increases tDR^-Gln^ levels in β- and non-β-cells. **A**, *TRMT10A* was silenced in clonal human β-cells EndoC-βH1 using one of our previously validated siRNAs (si*TRMT10A*). *TRMT10A* mRNA expression was normalized to *ACTB* and expressed as *2^-^*^Δ*Ct*^. **B** and **E**, schematic representation of tDR^-1:18-Gln-CTG-1-M10^ and tDR^-1:29-Gln-CTG-1-M3^ on top of their parental tRNA. **C, D, F**, and **G**, RT-qPCR data showing increased tDR^-1:18-Gln^ and tDR^-1:29-Gln^ levels in *TRMT10A-*silenced EndoC-βH1 cells (**C** and **F)**, and *TRMT10A* KO St7 islets (**D** and **G)**. Data was normalized to hsa-Let7-7F-5p and is expressed as 2^-ΔCt^. **H** and **I**, Levels of the parental tRNA (tRNA^Gln-CTG^) in control and *TRMT10A* KO cells assessed by RT-qPCR (**I**). The position of primers used for the amplification is shown in the scheme (**H**). Data was normalized to the snRNA U6 and is expressed as 2^-ΔC^. **J**, Schematic representation of the LI-FISH assay designed to detect tDR^-1:29-Gln^. **K**, Representative confocal images showing increased tDR^-1:29-Gln^ levels in the cytoplasm of insulin positive and insulin negative *TRMT10A* KO cells. No-ligase is a negative control of the essay. All pictures were taken using equal exposition settings and magnification. Scale bar, 10 µm. **L**, Subcellular fractionation of iPSC-derived St7 *TRMT10A* KO aggregates followed by RT-qPCR. 5S rRNA and U6 expression were used as cytoplasmic (CYT) and nuclear (NUC) controls, respectively. 5S, U6, tDR^-1:18-Gln^ and tDR^-1:29-Gln^ levels are expressed as % in each compartment. Subcellular fractionation data are means ± SE of n=2 independent differentiations. Panel **J** was created with BioRender (https://BioRender.com).

### tDR^-1:29-Gln^ interacts with and lowers hnRNPM expression in pancreatic β-like cells

tDRs modulate biological processes through different mechanisms, including protein interactions(Bayazit et al., 2022; Brozzi et al., 2024; Jacovetti et al., 2024; Kuhle et al., 2023). To assess putative tDR^-1:29-Gln^ interactions we performed an exploratory RNA-pull down assay in EndoC-βH1 cells using a 3’-biotinylated synthetic tDR^-1:29-Gln^ mimic or a scrambled 3’-biotinylated RNA control(Bayazit et al., 2022; Jacovetti et al., 2024), and identified bound proteins by mass spectrometry (Fig. 6A). After peptide identification using MSFragger, candidate interacting proteins were shortlisted based on the following criteria: (i) detection of at least 15 unique peptides for a given protein in the tDR^-1:29-Gln^ condition, and (ii) absence of peptides for that protein in the scramble RNA condition. This yielded 65 proteins potentially associated with tDR^-1:29-Gln^. KEGG pathway analysis revealed enrichment in splicing-associated proteins (Fig. 6B and Supplementary Fig. 9). Heterogeneous nuclear ribonucleoprotein M (hnRNPM), a multifunctional RNA-binding protein crucial for alternative splicing, gene expression regulation and cellular stress responses(Asano et al., 1991; Lv et al., 2025; West et al., 2019; Zheng et al., 2024), emerged as the top hit, with 36 unique peptides and 102 total peptides. We validated the tDR^-1:29-Gln^-hnRNPM association by Western blot in independent RNA pull-down assays (Fig. 6C) and by confocal microscopy using Li-FISH in *TRMT10A* KO St7 islets (Fig. 6D). Previous studies in non-β-cells indicate that, under steady state conditions, hnRNPM is primarily localized in nuclear structures, such as paraspeckles and the nuclear matrix, although it can translocate to the cytoplasm in disease or in response to cellular signals(Geuens et al., 2016). hnRNPM was detected in the nucleus and cytoplasm of *TRMT10A* KO β-like and non- β-like cells; its colocalization with tDR^-1:29-Gln^ occurred mainly in the perinuclear region (Fig. 6D).

**Fig. 6.**
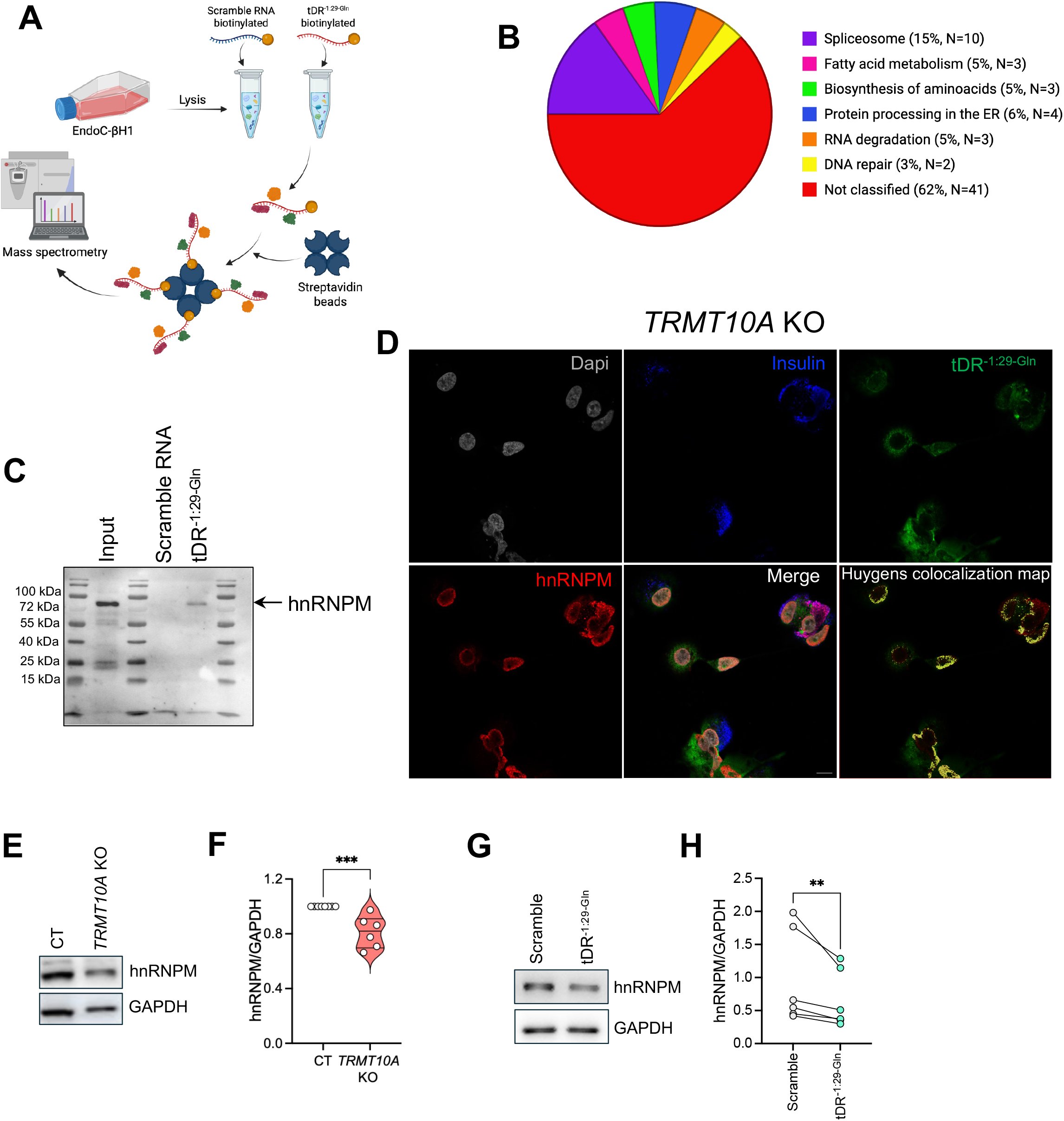
tDR^-1:29-Gln^ interacts with hnRNPM. **A**, Schematic representation of the exploratory RNA pull-down assay performed to identify proteins associated with tDR^-1:29-Gln^ in EndoC-βH1 cells. **B**, KEGG pathway analysis performed with the proteins potentially associated with tDR^-1:29-Gln^. **C**, Western blot of a representative independent RNA pull down assay confirming the interaction of tDR^-1:29-Gln^ with hnRNPM in EndoC-βH1 cells. **D**, Representative confocal images of *TRMT10A* KO St7 islets showing hnRNPM (red), insulin (blue) and tDR^-1:29-Gln^ (detected by LI-FISH,green). Nuclei were stained with Dapi (grey). Huygens analysis showed a partial co-localization between tDR^-1:29-Gln^ and hnRNPM (Huygens colocalization map). Images acquired at 63X magnification. Scale bar, 10 µm. **E-H**, hnRNPM protein expression assessed by Western blot in **E-H**, hnRNPM protein expression in isogenic control (CT) and *TRMT10A* KO St7 islets (**E-F**), and EndoC-βH1 cells transfected with a scramble RNA or synthetic tDR^-1:29-Gln^ mimic (**G-H**). **E** and **G**, Representative Western blot images of n=6 independent experiments. **F** and **H**, Quantification of the blots. ** p<0.01, *** p<0.001 by unpaired t-test (**F**) or ratio paired t-test (**H**). Panels **A** and **B** were created with BioRender (https://BioRender.com).

Given that tDRs can alter the function or the stability of their target proteins(Yu et al., 2021), we next examined whether hnRNPM expression was altered in *TRMT10A* deficiency. *TRMT10A* KO St7 islet-like aggregates showed reduced hnRNPM protein expression by Western blot (Fig. 6E-F). Furthermore, transfection of EndoC-βH1 cells with synthetic tDR^-1:29-Gln^ mimics replicated the reduction of hnRNPM protein abundance (by 30%, Fig. 6G-H). In sum, these results suggest that tDR^-1:29-Gln^ interact with hnRNPM in *TRMT10A*-deficient cells and lowers its expression.

### hnRNPM interacts with PTBP1 in pancreatic β-cells and controls insulin content and processing

STRING analysis indicated that hnRNPM potentially interacts with several hnRNP family members including hnRNPI (also known as polypyrimidine tract-binding protein, PTBP1) (Fig. 7A). This interaction was experimentally validated in 293FT cells(Hu et al., 2020) and mouse testis(Lv et al., 2025). Considering that PTBP1 plays an important role in β-cells by regulating insulin mRNA stability and translation and *PCSK1* expression(Fred and Welsh, 2009; Han et al., 2012; Jeong et al., 2018; Magro and Solimena, 2013), we investigated whether hnRNPM and PTBP1 interact in human β-cells. To this end, we used PTBP1 as bait in a co-immunoprecipitation assay in EndoC-βH1 cultured at normal or high glucose concentrations. Interestingly, hnRNPM and PTBP1 interacted, especially under stimulatory glucose conditions (Fig. 7B-C). To assess the functional role of hnRNPM in β-cells, we silenced its expression in EndoC- βH1 cells (Fig. 7D). Similar to the phenotype observed in *TRMT10A*-deficient cells, *hnRNPM* knockdown significantly reduced *PCSK1* mRNA and protein expression (Fig. 7E-G). Moreover, *hnRNPM*-silenced cells displayed lower insulin content (Fig. 7H), unchanged proinsulin levels (Fig. 7I) and a significantly increased proinsulin to insulin ratio (Fig. 7J) indicative of impaired proinsulin processing. PTBP1 protein expression remained unchanged upon hnRNPM silencing (Supplementary Fig. 10). Given that transfection of tDR^-1:29-Gln^ mimics reduced *hnRNPM* levels (Figure 6G-H), we next examined their impact on *PCSK1* expression. tDR^-1:29-Gln^ mimics transfection reduced *PCSK1* mRNA expression in EndoC-βH1 cells (Fig. 7K) supporting a potential contribution of these fragments to the impaired insulin processing observed in *TRMT10A*-deficient cells. Collectively, these findings identify a previously unrecognized interaction between hnRNPM and PTBP1 in human β-cells and suggest that hnRNPM contributes to the regulation of *PCSK1* expression and proinsulin maturation. Furthermore, our data are consistent with a model in which *TRMT10A* deficiency-induced tDR^-1:29-Gln^ fragments impair *PCSK1* expression, at least in part, through the reduction of hnRNPM abundance and/or activity. This effect may alter the function of the hnRNPM–PTBP1 regulatory complex without affecting PTBP1 expression itself, although the precise mechanism warrants further investigation (Fig. 7L).

**Fig. 7.**
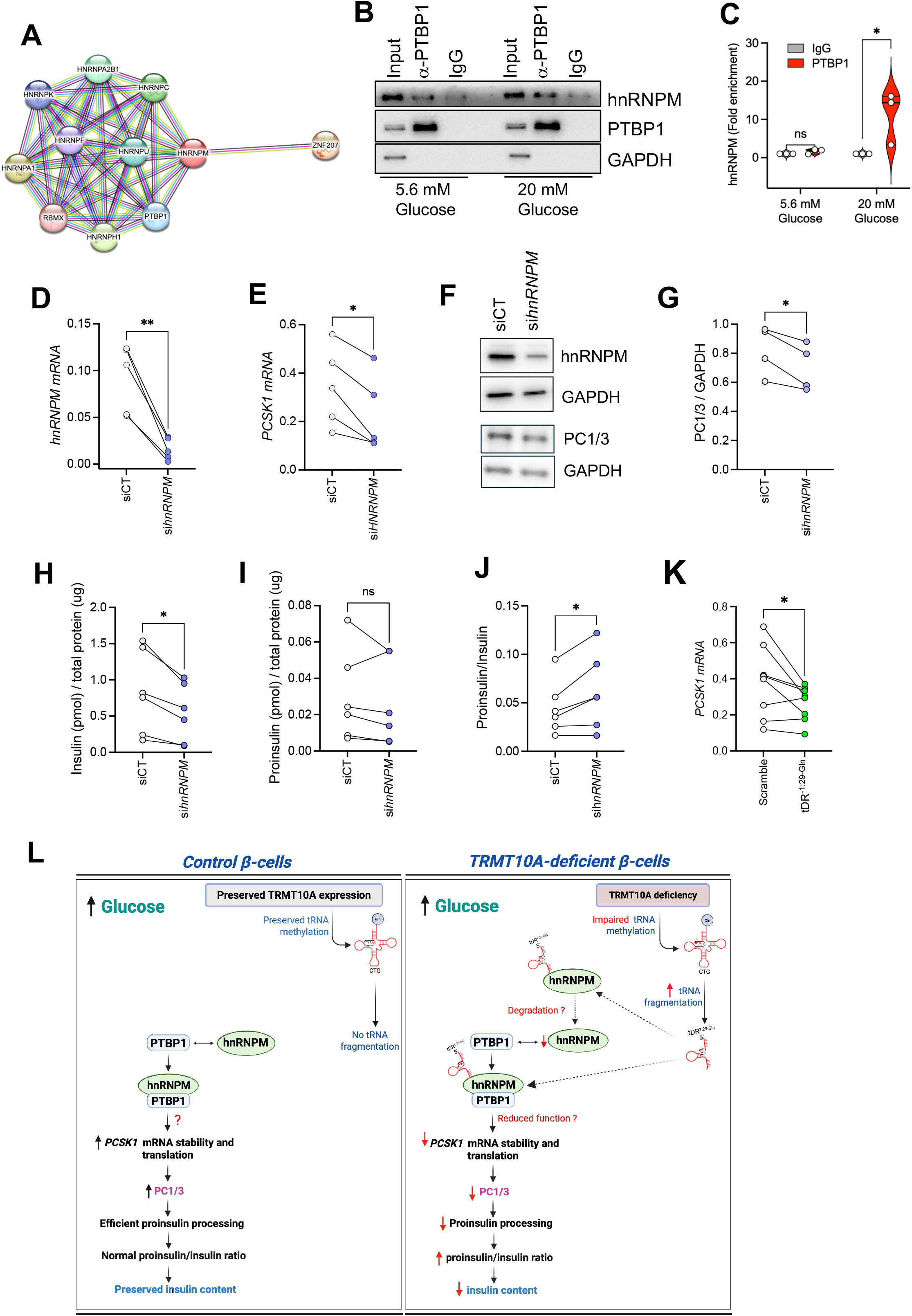
hnRNPM regulates PCSK1 expression and insulin processing. **A**, Protein-protein interaction network generated using STRING https://string-db.org/ showing predicted interactions between hnRNPM and other hnRP family members including PTBP1. **B-C**, Co-immunoprecipitation validating the interaction between PTBP1 (bait) and hnRNPM in EndoC-βH1 cells exposed to 5.6 mM glucose or 20 mM glucose for 16h. Mouse IgG served as negative control. **B**, Representative image of n=3 independent experiments. **C**, Quantification of hnRNPM protein bound to PTBP1 or to mouse IgG relative to the input. Values were normalized to the IgG control and expressed as fold enrichment. **D-J**, *hnRNPM* silencing in EndoC-βH1 cells. **D** and **E**, *hnRNPM* and *PCSK1* mRNA expression assessed by RT-qPCR. Data was normalized to the geometric mean of *ACTB* and *VAPA* and is expressed as 2^-ΔCt^. **F** and **G**, PC1/3 protein levels in *hnRNPM*-silenced EndoC-βH1 cells. **F**, Representative Western blot of n=4 independent experiments. **G**, Quantification of the blots. **H**, Insulin and **I**, pro-insulin content of *hnRNPM*-silenced EndoC-βH1 cells measured by ELISA. **J**, Proinsulin to insulin ratio. **K**. PCSK1 mRNA expression in EndoC-βH1 cells transfected with a scramble RNA or the synthetic tDR^-1:29-Gln^ mimics. **L**, Scheme illustrating the proposed role of hnRNPM-PTBP1 complexes in control and *TRMT10A*-deficient human β-cells. In *TRMT10A*-competent β-cells, glucose stimulation promotes the interaction between hnRNPM and PTBP1, potentially contributing to PCSK1 mRNA stabilization and enhanced translation. In contrast, under *TRMT10A*-deficient conditions, increased levels of tDR^-1:29-Gln^ reduce hnRNPM abundance and/or function potentially affecting hnRNPM-PTBP1 complexes and PCSK1 expression. Symbols represent independent experiments. * p<0.05, ** p<0.01, ns: not significant, by ratio paired t-test. Panel K was created with BioRender (https://BioRender.com).

## Discussion

In this study we show that *TRMT10A* deficiency disrupts β-cell development and function, leading to impaired proinsulin processing, reduced insulin content and secretion, heightened oxidative stress, and broad transcriptional dysregulation. In addition to these global cellular effects, our findings suggest that *TRMT10A-*dependent tRNA^Gln^ fragmentation contributes to β-cell dysfunction via its interaction with hnRNPM potentially affecting hnRNPM-PTBP1-mediated regulation of *PCSK1* expression and proinsulin processing.

The developmental defects of *TRMT10A*-deficient β-like cells arose at pancreatic endoderm stage, and were marked by reduced NKX6.1 expression, a transcription factor essential for β-cell development, identity and proliferation(Aigha and Abdelalim, 2020; Schaffer et al., 2013). During pancreas development, three NKX6.1^+^ cell populations have been described: *PDX1^+^/NKX6.1^+^* pancreatic progenitors, *NGN3^+^/NKX6.1^+^* endocrine progenitors, and *INS^+^/PDX1^+^/NKX6.1^+^* β-cells(Aigha and Abdelalim, 2020). In mice, *Nkx6.1* and *Pdx1* co-expression is critical for committing pancreatic progenitors to the monohormonal β-cell lineage. Pancreatic progenitors expressing *Pdx1* but lacking *Nkx6.1,* as in the *TRMT10A*-deficiency scenario, generate β-cells with reduced insulin content, impaired glucose uptake and insulin secretion, and increased polyhormonal cells(Aigha and Abdelalim, 2020; Taylor et al., 2013). Although *TRMT10A*-deficient β-like cells exhibited reduced insulin content and secretory function, we did not observe an increase in polyhormonal cells, which may reflect residual NKX6.1 in our model or species-specific differences between human and murine pancreas development. The sustained reduction in *NKX6.1* likely underlies the lower proportion of β-cells in *TRMT10A*-deficient islet-like aggregates compared to control lines. *PAX4* expression was also reduced in *TRMT10A*-deficient St7 aggregates. While *Pax4* plays a crucial role in mouse β-cell specification during pancreas development(Sosa-Pineda et al., 1997), it is dispensable for human β-cell differentiation but important for β-cell maturation(Lau et al., 2023). *PAX4* silencing in EndoC-βH1 cells reduced insulin secretion and content(Lau et al., 2023), and *PAX4* KO iPSC-derived islet-like aggregates had downregulated β-cell maturation and insulin secretion-related genes (*MAFA, ISL1, SLC17A6, PCSK2*), derepression of α-cell genes (*ARX*, *GCG* and *TTR*) and increased polyhormonal cells(Lau et al., 2023). While *TRMT10A* KO islet-like aggregates exhibited impaired insulin secretion and content, there was no increase in α-cell number or genes (*GCG, ARX and TTR)*, suggesting that *PAX4* inhibition does not mediate the phenotype.

Beyond islet-development genes, *TRMT10A* deficiency also resulted in dysregulation of genes associated with neurological and neurodevelopmental phenotypes that parallel patient manifestations, including intellectual disability, epilepsy, and microcephaly. Genes involved in synaptic vesicle cycling, neurotransmission, neuronal excitability and cortical development such as *CACNA1E, NSF*, *NPTX2*, and others identified in our transcriptomic datasets were significantly altered. These transcriptional changes mirror the clinical spectrum reported in *TRMT10A*-deficient individuals and highlight the broader impact of impaired tRNA modification on neuroendocrine systems.

*ABCC8* expression was also reduced in *TRMT10A*-deficient cells, consistent with the hyperinsulinemic hypoglycemic episodes reported in some *TRMT10A*-deficient individuals(Gillis et al., 2014; Siklar et al., 2021; Zung et al., 2015). Dominant inactivating mutations of *ABCC8* are associated with a broad clinical spectrum, ranging from asymptomatic to transient or persistent hyperinsulinemic hypoglycemia that can be treated with diazoxide(De Franco et al., 2020; Kapoor et al., 2011). We did not, however, observe signs of insulin hypersecretion in *TRMT10A*-deficient β-like cells, and the response to gliclazide and diazoxide was preserved, suggesting that the K_ATP_ channel functions normally.

*TRMT10A*-deficient β-like cells also showed blunted insulin secretion in response to glucose but not KCl. Chronic oxidative stress may impair glucose responsiveness without affecting KCl-induced membrane depolarization and Ca²⁺ influx(Stancill and Corbett, 2021). In line with our previous published findings in *TRMT10*A-silenced INS-1E and EndoC-βH1 cells(Cosentino et al., 2018), *TRMT10A* KO islets exhibited signs of oxidative stress, likely due to lesser antioxidant capacity. Consistent with this, Seahorse profiling revealed diminished spare respiratory capacity and proton leak, together with reduced FCCP-stimulated respiration and increased ATP production- features indicative of a rigid mitochondrial state prone to ROS generation. In addition, NAD(P)H autofluorescence was markedly lower in *TRMT10A*-deficient aggregates, indicating diminished reducing-equivalent availability and further compromising the cells’ ability to buffer oxidative stress. Besides *UCP2*, there was downregulation of catalase (*CAT*), which detoxifies H_2_O_2_; glutathione peroxidase 7 (*GPX7*), an ER-resident enzyme that scavenges H_2_O_2_ generated during oxidative protein folding(Pei et al., 2023); and peroxiredoxins 3 and 5 (*PRDX3* and 5) which primarily protect mitochondria from oxidative stress(Wolf et al., 2010) (Supplementary Fig. 5). It was shown that low *PRDX3* expression impairs insulin secretion and increases stress-induced apoptosis(Wolf et al., 2010), two features of *TRMT10A*-deficient β-cells. When considered alongside the reduced spare respiratory capacity and markedly lower NAD(P)H levels, this coordinated loss of antioxidant defenses offers a mechanistic basis for the oxidative stress previously reported in *TRMT10A*-deficient β-cells(Cosentino et al., 2018).

Insulin secretion by *TRMT10A*-deficient LTC β-like cells was also impaired in response to exendin-4, a GLP-1 receptor agonist. This defect is consistent with the twofold reduction in *GLP1R* expression, which likely dampens the incretin response. Activation of the GLP-1R in β-cells enhances insulin biosynthesis, potentiates glucose-stimulated insulin secretion, and exerts longer-term trophic effects on β-cell survival, proliferation, and neogenesis(Marzook et al., 2021).

The impaired insulin secretion observed in *TRMT10A*-deficient β-like cells is closely linked to a defect in proinsulin processing and maturation. This processing deficit likely stems from a marked reduction in *PCSK1* (PC1/3), the principal protease responsible for proinsulin cleavage in human β-cells, with additional decreases in *PCSK2*, *CPE*, and *ERO1B* further compromising the maturation process. Consistent with our findings, PC1/3 deficiency has been shown to impair proinsulin processing in human β-cells(Meier et al., 2022) and *WFS1*-deficient iPSC-derived β-like cells(Torchio et al., 2025). Moreover, reduced PC1/3 and CPE levels have similarly been reported in islets from individuals with type 1 diabetes(Sims et al., 2019a; Sims et al., 2019b). Together, the loss of these proinsulin-processing components results in markedly diminished insulin content, providing a mechanistic explanation for the profound secretory defects of *TRMT10A*-deficient cells.

Furthermore, several pathways associated with ER-to-Golgi vesicle-mediated transport, Golgi organization, and vesicle exocytosis were significantly downregulated, as indicated by both RNA-seq and proteomic datasets. Efficient ER-Golgi trafficking is essential for the proper folding, quality control, and forward transport of proinsulin, as well as for the acidification and maturation of secretory granules(Boyer et al., 2023). Consistent with this, *TRMT10A*-deficient cells exhibited reduced expression of *ATP6AP1*, a V-ATPase accessory subunit required for granule acidification, which is a key step in enabling PC1/3 catalytic activity(Zavarzadeh et al., 2025). Downregulation of *ERO1B*, an ER-resident oxidoreductase involved in disulfide bond formation during proinsulin folding(Awazawa et al., 2014) further supports compromised function of the early secretory pathway. Together, these defects in the ER-Golgi network likely synergize with the loss of PC1/3 and related processing enzymes to impair proinsulin maturation, reduce insulin content, and exacerbate β-cell dysfunction. While the precise origin of these trafficking abnormalities remains unclear, they may reflect a combination of altered transcriptional programs, increased oxidative load, and disrupted cellular homeostasis resulting from *TRMT10A* deficiency. Our previous work(Cosentino et al., 2018) and current data demonstrate that *TRMT10A* deficiency leads to tRNA fragmentation. Cytoplasmic tDR^Gln-CTG^ levels were increased in *TRMT10A-*deficient β- and non-β-cells without changes in parental tRNA^Gln-CTG^ level, this paper, or aminoacylation(Cosentino et al., 2018). This suggests that the hypomodification and fragmentation of tRNA^Gln^ in human islets does not compromise its availability for protein synthesis. Indeed, the RedRibbon overlap between *TRMT10A*-deficient islet-like transcriptome and proteome does not point to major translational defects. A very different scenario was reported in *trmt10a* null mice, where the full tRNA^Gln-CTG^ and tRNA initiator methionine levels were reduced in various tissues in the absence of tRNA fragmentation(Tresky et al., 2024). This impaired translation of corresponding codons in neuronal mRNAs. The mice showed impaired hypothalamic synaptic plasticity and memory but, unlike *TRMT10A* deficient patients, did not develop microcephaly or diabetes(Tresky et al., 2024). The smaller brain size of *trmt10a* null mice was proportional to their overall reduced body size, and their glucose tolerance was preserved under chow and high fat diet(Tresky et al., 2024), indicating that *TRMT10A* deficiency displays major interspecies differences.

We previously showed that tRNA^Gln^ fragmentation contributes to *TRMT10A*-deficiency-mediated β-cell apoptosis(Cosentino et al., 2018). Here we demonstrate that tDR^-1:29-Gln^ may also regulate insulin processing through its interaction with hnRNPM-PTBP1. tDR^-1:29-Gln^ mildly reduce hnRNPM protein levels, although interference with hnRNPM activity cannot be excluded. Other tDRs have been shown to interact with hnRNPM. Indeed, cleavage of tRNA^Gly-GCC^ into tDR^Gly-GCC^ (5′-tDR^-GlyGCC^) by IRE1α during ER stress was shown to promote cancer cell proliferation and survival through interactions with hnRNPM and hnRNPH2 across species(Jin et al., 2024). Such fragments may play an evolutionarily conserved role in cellular adaptation to ER stress(Jin et al., 2024). hnRNPM is a multifunctional RNA-binding protein essential for alternative splicing, gene expression regulation and cellular stress responses. It also acts as an innate immunity modulator of antiviral activity(Asano et al., 1991; Cao et al., 2019; Lv et al., 2025; West et al., 2019; Zheng et al., 2024; Zhong et al., 2023). Interestingly, the *TRMT10A*-deficient islet-like omics revealed downregulated antiviral innate immune pathways and negative regulation of type I interferon signaling. Whether these changes result from tDR^-1:29-Gln^-mediated hnRNPM inactivation remains to be investigated.

PTBP1 is a ubiquitous RNA-binding protein that plays an important role in β-cells by controlling proinsulin mRNA stability and translation, as well as translation of other secretory granule proteins via a cap-independent mechanism involving PTBP1 binding to the 5’-UTR of target mRNAs(Magro and Solimena, 2013; Tillmar et al., 2002). PTBP1 functions through interaction with other proteins, including hnRNPC, hnRNPK and hnRNPE, and changes in its subcellular localization. Upon glucose stimulation, PTBP1 translocates from the nucleus to the cytosol where it modulates mRNA stability and/or translation(Magro and Solimena, 2013). Interestingly, it was shown that PTBP1 silencing reduces secretory granule stores and expression of secretory granules components including PC1/3(Knoch et al., 2004). Here we demonstrate that hnRNPM interacts with PTBP1 in EndoC-βH1 cells, primarily under high glucose conditions. In 293FT cells(Hu et al., 2020) and mouse germ cells(Lv et al., 2025) this interaction regulates alternative splicing and male fertility. We show that hnRNPM silencing in human β-cells impairs *PCSK1* (PC1/3) expression, proinsulin processing and insulin content, suggesting that hnRNPM supports PTBP1 function on key mRNA targets in pancreatic β-cells. In particular, our data suggests that, in control human β-cells, hnRNPM and PTBP1 interact upon glucose stimulation to potentially stabilize *PCSK1* mRNA and promote its translation. In *TRMT10A*-deficient β-cells, tDR^-1:29-Gln^ binds hnRNPM and may perturb hnRNPM–PTBP1 interactions, contributing to reduced *PCSK1* (PC1/3) expression and impaired proinsulin processing (Fig. 7L).

tRNA fragmentation has been associated with diabetes and its complications, and Parkinson’s disease, Alzheimer’s disease and multiple sclerosis(Bayazit et al., 2022; Brozzi et al., 2024; Cosentino et al., 2018; Jacovetti et al., 2024; Winek and Soreq, 2025; Zhang et al., 2025). In pancreatic islets and brain, some tDRs contribute to tissue homeostasis whereas others promote disease(Bayazit et al., 2022; Brozzi et al., 2024; Jacovetti et al., 2024; Winek and Soreq, 2025). For example, tRNA halves from tRNA^His^ and tRNA^Glu^ are abundant in neonatal rat islets and support β-cell-mass expansion and insulin secretory capacity(Bayazit et al., 2022). tDRs from mt-tRNA^Leu-TAA^, which are reduced in type 2 diabetes models, interact with electron transfer chain complexes to regulate mitochondrial function and insulin secretion(Jacovetti et al., 2024). Brozzi et al demonstrated that in type 1 diabetes models the tDR pool of mouse β-cells and immune cells is altered by proinflammatory cytokines, and that the transfer of tDRs from immune cells to β-cells sensitizes β-cells to apoptosis (Brozzi et al., 2024). Given their wide-ranging influence on β-cell biology, the potential impact of tDRs on proinsulin processing and β-cell function warrants further investigation. While our data establish a robust association between *TRMT10A* deficiency and altered *NKX6.1* expression, they do not directly resolve the underlying molecular mechanisms. It is possible that additional tDRs are generated under *TRMT10A*-deficient conditions in human β-cells contributing to the observed phenotype, or that increased m^6^A RNA methylation, as shown in *TRMT10A*-deficient HEK293 cells(Ontiveros et al., 2020), negatively impact β-cell development. Dissecting the relative contribution of these pathways will be an important focus of future studies.

In conclusion, our study identifies TRMT10A as a critical regulator of human β-cell development and function.

Loss of this tRNA methyltransferase leads to the accumulation of tDR^-1:29-Gln^, which is associated with impaired *PCSK1* mRNA stability and translation, defective proinsulin processing, and marked β-cell dysfunction. Our findings support a model in which tDR^-1:29-Gln^ could perturb hnRNPM–PTBP1 interactions, thereby contributing to these defects.

Moreover, the neuronal expression signatures in our omics analyses suggest that similar unappreciated regulatory mechanisms may affect human neuronal development and function in TRMT10A patients.

Collectively, our findings position TRMT10A as a key determinant of β-cell health and reveal molecular pathways that could be targeted to prevent β-cell failure in monogenic and polygenic diabetes.

## Acknowledgements

We thank Anyishaï Musuaya, Angeline Bilheu, Jessica Capitaine and Isabelle Millard for expert technical assistance.

## Author contributions

C.M.C., M.N.A. and K.B. contributed equally to this work. M.N.A., B.J.S., and J.G. generated the *TRMT10A* KO iPSC-cells and K.B. the *TRMT10A*-corrected control iPSC-cells. T.S. and M.C. implemented long-term iPSC-β-cell differentiation. B.S. and M.N.A. developed the Li-FISH assay. S.S., K.B., and J.C.J. performed calcium and NADPH measurements. T.S. and K.B. performed the dynamic insulin secretion studies. D.C. and V.I. did the mass spectrometry analysis in *TRMT10A* KO St7 islets. K.G. and J.P. performed the mass spectrometry analysis in *TRMT10A*-silenced β- cells. C.C. and R.R. performed RNA sequencing. C.M.C., K.B., N.P. and A.O.G. performed iPSC differentiation and quality controls. C.M.C. performed the confocal analysis. C.M.C., M.N.A., K.B., A.O.G., B.J. S., T.S., C.C., S.S. J.C.J. M.C. and M.I.E. designed the experiments, collected and interpreted the data. M.I.E. performed the bioinformatic analyses, planned the project, and supervised the study. M.I.E. and M.C. wrote the manuscript. All authors read, corrected, and approved the manuscript. M.I.E. is the warrantor of this work.

## Data, and materials availability

All data necessary to evaluate and reproduce the findings of this study are provided within the main text and the Supplementary Materials. The datasets generated and analyzed during this work have been deposited in the Gene Expression Omnibus and the PRIDE repositories and will be publicly available upon acceptance of the manuscript (private access numbers will be provided to the reviewers under editorial request). This study did not generate any new materials.

## Materials and Methods

*Detailed methods are provided as Supplementary Materials*

### Ethical approval

The *TRMT10A*-mutant iPSC line HEL122.2 was reprogrammed from adult skin fibroblasts(Cosentino et al., 2018). The human iPSC line 1023A was kindly provided by Dieter M Egli, University of Columbia. iPSC culture, gene editing and differentiation into human islet-like aggregates cells was approved by the Ethical Committee of the Erasmus Hospital, Université Libre de Bruxelles, reference P2019/498 and A2024/211.

### CRISPR–Cas9 Gene Editing of iPSCs

*TRMT10A* knockout (KO) and *TRMT10A* corrected iPSCs were generated by CRISPR-Cas9-mediated gene editing of the validated control line 1023A, and the *TRMT10A* mutant (p.Arg127*) patient-derived HEL122.2 iPSC line, respectively. The gRNAs and donor sequence were designed and synthesized at Integrated DNA Technologies (IDT), sequences are provided in Supplementary Table 1. Potential off-target sites (Supplementary Table 2) in the *TRMT10A*-edited clones and their parental iPSC lines were PCR-amplified and sequenced. Primer sequences are listed in Supplementary Table 3. One homozygous KO line and two corrected clones were selected. Off-target sites predicted *in silico* were PCR-amplified and sequenced. Chromosomal integrity was verified by KaryoStat™ or microarray/iCS-digital™ PSC assay. Pluripotency and tri-lineage potential were confirmed by marker expression and embryoid body differentiation. Antibodies used are listed in Supplementary Table 4.

### Cell Culture

EndoC-βH1 cells were grown on ECM + fibronectin-coated plates in complete EndoC-βH1 medium (Supplementary Table 5) at the cell densities specified in Supplementary Table 6.

Human iPSCs were differentiated into islet-like aggregates using a previously reported seven-stage protocol (Virgilio et al., 2025). Stage-7 (St7) aggregates were maintained in suspension in St-7 medium (Supplementary Table 7) in ultra-low attachment plates under rotation and matured a further month in long-term culture medium (Supplementary Table 8) to enhance β-cell functionality (Virgilio et al., 2025). Differentiation efficiency was assessed by qPCR and immunostaining. The antibodies and primers used are listed in Supplementary Tables 4 and 9, respectively.

### Dispersion of iPSC-derived Islets

St7 or long-term aggregates were incubated in Accumax for 8–10 min at 37 °C, gently triturated, and replated on Matrigel with ROCK inhibitor. After recovery, cells were switched to complete human islet medium (Supplementary Table 10) for 24 h prior to treatment.

### RNA Interference

EndoC-βH1 cells were transfected overnight with 30 nM siRNAs targeting *TRMT10A*, *hnRNPM*, or a non-targeting control using Lipofectamine RNAiMAX. Medium was replaced after 16 h and cells processed 96 h post-transfection. Transfection medium composition, lipofectamine concentrations, and siRNA sequences are provided in Supplementary Tables 5, 6 and 11, respectively.

### Transfection with tDR^-1:29-Gln^ mimics

Cells were transfected with 60 nM tDR^-1:29-Gln^ mimics or scramble control using Lipofectamine 2000. Medium was changed after 16 h, and cells collected 48 h post-transfection. Lipofectamine concentration and RNA mimic sequences are provided in Supplementary tables 6 and 12, respectively.

### RNA Extraction, Reverse Transcription, and qPCR

Total RNA was extracted using Direct-zol kits. Reverse transcription was performed from 400 ng RNA using random primers, and qPCR was performed with SYBR Green chemistry on a MyiQ2 system. Expression was normalized to *ACTB* or *VAPA*. Primer sequences are listed in Supplementary Table 9.

### Quantification of tDR^-1:29-Gln^, tDR^-1:18-Gln^ and full tRNA^-Gln-CTG^ by RT-qPCR

Total RNA was extracted as described above, and prior to reverse transcription it was demethylated with AlkB/AlkB-D135S and re-purified. Full-length tRNA was reverse-transcribed using RevertAid, while tDRs were measured with miRCURY LNA assays. U6 or hsa-let-7f-5p served as endogenous controls. Relative abundance was calculated using the ΔCt method. The primers used are provided in Supplementary Table 9 and sequences of the analyzed tDRs in Supplementary Table 13.

### Ligation-Fluorescent In Situ Hybridization (LI-FISH)

EndoC-βH1 cells and dispersed iPSC-derived islet-like aggregates were fixed with 4% paraformaldehyde and permeabilized with 0.5% Triton X-100. A tDR^-1:29-Gln^-specific stem-loop was hybridized and ligated usingT4 RNA ligase. A digoxigenin-labelled probe was then hybridized to the ligation product and detected using a mouse anti-digoxigenin antibody followed by anti-mouse-fluorescent antibody. Images were taken using Zeiss LSM 780 or Axio Observer microscopes. Analyses were done with ImageJ and Huygens software. A “no-ligase” control was used to verify the signal specificity. Antibodies and dilutions used are listed in Supplementary Table 4.

### Subcellular Fractionation

Cytoplasmic and nuclear fractions were isolated using the PARIS kit. Fraction purity was confirmed with 5S rRNA and U6 snRNA. Demethylated RNA from each fraction was analyzed for tDR^-1:29-Gln^ and tDR^-1:18-Gln^ by RT-qPCR.

### Western Blotting

Islet-like aggregates or EndoC-βH1 cells were lysed in Laemmli buffer, separated by SDS-PAGE, and transferred to PVDF membranes. Blots were probed with primary antibodies overnight, washed, and incubated with HRP-conjugated secondaries. Signals were detected by chemiluminescence and quantified using Fiji. Antibodies and dilutions used are listed in Supplementary Table 4.

### Co-Immunoprecipitation

EndoC-βH1 lysates (standard or high-glucose conditions) were prepared in T7 co-immunoprecipitation buffer (20mM HEPES pH 7.9, 150mM NaCl, 5mM EDTA, 1% NP-40, 1mM DTT, 10% Glycerol) with RNase and protease inhibitors. After pre-clearing, lysates were incubated with anti-PTBP1 or control IgG, followed by Dynabeads capture and extensive washing. Eluted proteins were examined by Western blot to assess hnRNPM–PTBP1 complex formation.

### Insulin Secretion Assays

#### Static non-sequential insulin secretion

St7 islet-like aggregates were pre-incubated in glucose-free Krebs buffer, then stimulated with low or high glucose, with or without diazoxide, gliclazide, or KCl. Supernatants were collected after 1 h for insulin quantification by ELISA.

#### Dynamic Insulin Secretion

Long-term matured aggregates were perifused sequentially with low glucose, high glucose, high glucose + Exendin-4, low glucose, and KCl. Fractions were collected every 4 min for insulin assessment by ELISA

#### Insulin and Proinsulin Content

Islet-like aggregates were lysed by sonication, and insulin/proinsulin content was measured by ELISA.

### NAD(P)H Autofluorescence

Islet-like aggregates were perifused with Krebs buffer in a heated chamber on the stage of an inverted microscope, and NAD(P)H autofluorescence (λ_ex 360 nm/λ_em 470 nm) was recorded every 10 s. After baseline acquisition, FCCP, diamide, and Kp-372 were applied to probe mitochondrial and redox responses.

### Seahorse assay

Dispersed St7 islet-like aggregates were plated in Seahorse miniplates and equilibrated in Krebs buffer. Basal and stimulated oxygen consumption rate (OCR) was recorded following injections of glucose, oligomycin, FCCP, and rotenone/antimycin. OCR values were normalized and used to compute ATP production, maximal respiration, proton leak, and spare respiratory capacity. OCR data was normalized to the last basal reading in each sample, considered as 1, and used to calculate different parameters associated with mitochondrial function as follows: *Non mitochondrial oxygen consumption*= Last OCR measured; *Maximal respiration*=First OCR measured after FCCP injection – non mitochondrial oxygen consumption; *H+ (Proton) leak*= Last OCR measured after Oligomycin injection – non mitochondrial oxygen consumption; *ATP production rate*= last OCR of basal measurement – Last OCR after Oligomycin injection; *Spare respiratory capacity*= Maximal respiration – basal respiration; *Acute response*=Last OCR after glucose injection – last OCR of basal measurement.

### Immunofluorescence

Cells were fixed with 4% paraformaldehyde, permeabilized with 0.5% Triton X-100 and blocked prior to overnight incubation with primary antibodies. After washing, secondary antibodies and Hoechst were applied for 1 h. Samples were mounted and imaged using fluorescence microscopy. Antibodies and dilutions used are listed in Supplementary Table 4.

### RNA pull-down and Mass Spectrometry

Potential protein partners of tDR^-1:29-Gln^ were identified using an exploratory RNA pull down assay followed by mass spectrometry, as previously described (Bayazit et al., 2022). 5’-biotinylated scramble RNA, or a 5’-biotinylated oligonucleotide mimicking the tDR-^1:29-Gln^ were denatured at 95 °C for 2 min, and refolded. Refolded biotinylated oligonucleotides were then incubated for 1h at RT with EndoC-βH1 lysates. Complexes were captured with streptavidin beads, washed, and processed for proteomics.

### Sample preparation for Mass Spectrometry analysis

Proteins from the RNA Pull down assay were processed directly on beads by resuspension in 8M Urea/50mM Tris-HCl (pH8.5).

Proteins from iPSC-derived St7 aggregates (n=7 isogenic control and 7 *TRMT10A* KO per group) were extracted in urea buffer, homogenized, sonicated, clarified, and quantified.

Proteins were then reduced, alkylated, diluted, and digested with Lys-C and trypsin. Tryptic peptides were purified by solid-phase extraction and dried.

### Mass Spectrometry acquisition

RNA pull-down peptides were analyzed by nanoLC–DDA on a TripleTOF 5600 system. Peptides from iPSC-derived islet-like aggregates were acquired in SWATH-DIA mode using a variable-window scheme. Detailed chromatography and MS settings are provided in Supplementary Methods.

### Proteomics data analysis

RNA pull-down data were searched with MSFragger/FragPipe, with peptide and protein FDR thresholds at 1%. DIA datasets were analyzed with DIA-NN using a curated spectral library. Differential abundance testing used FragPipe-Analyst with missing-value and fold-change filters (adjusted p < 0.05; |log2FC| ≥ 0.58).

### Proteomics in *TRMT10A*-silenced EndoC-βH1 cells

*TRMT10A* expression was silenced by RNA interference in EndoC-βH1 cells (n=4 independet experiments) using two previously validated *TRMT10A* siRNAs (siTRMT10A#1 and #2)(Cosentino et al., 2018). Cells transfected with a control siRNA were used as control. siRNA sequences are provided in Supplementary Table 11. Lysates were reduced, alkylated, and digested with Lys-C and trypsin. Peptides were cleaned, fractionated by SCX, and analyzed by LC–MS/MS on a Q-Exactive HF. Data were searched with MaxQuant/Andromeda against UniProt.

### RNA Sequencing

RNA from isogenic control and *TRMT10A* KO St7 islet-like aggregates (n = 8 each) was isolated, quality-checked (RIN ≥ 7.6), and processed for TruSeq stranded RNA-seq. Reads were trimmed, aligned to GRCh38, and analyzed for differential expression FDR p < 0.05; |log2FC| ≥ 0.58). Enrichment analysis used DAVID, and plots were generated in SRplot.

### Statistical analysis

Statistical analyses were performed using GraphPad Prism version 10.5.0. For group data, results are presented as violin plots with individual datapoints. Paired data are shown as column charts with individual symbols connected by lines. Each data point represents an independent experiment. Non-normally distributed variables were log-transformed prior to statistical testing. Comparisons between two groups were performed using paired or unpaired t-test as suitable. Comparisons between three or more groups were analyzed using two-way ANOVA (or mixed effects analysis when missing values were present) followed by Sidak’s or Holm-Sidak correction for multiple comparisons, as recommended by the software. A p value or q value <0.05 was considered statistically significant.

---

**Supplementary Fig. 1.**
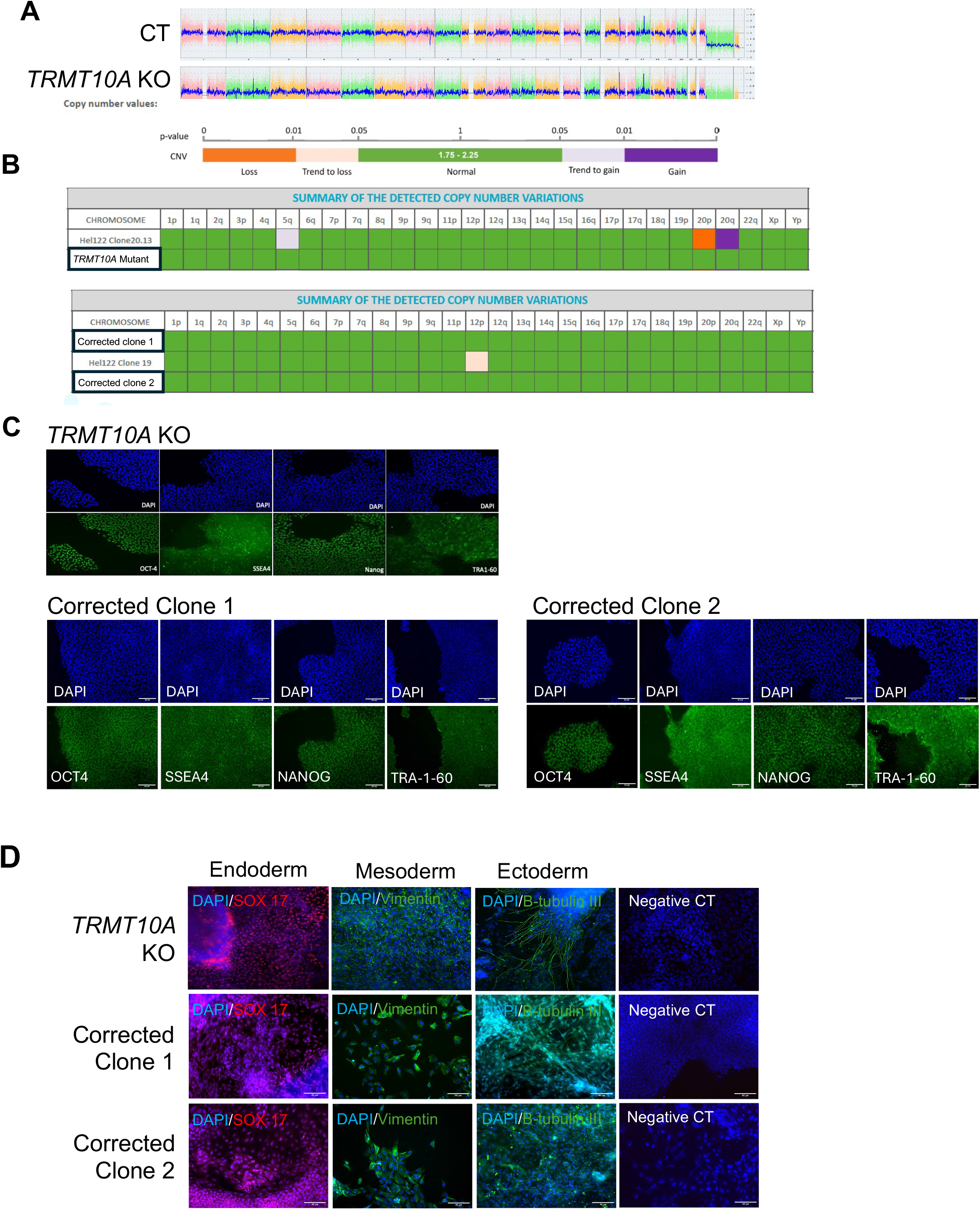
Quality control of TRMT10A KO and TRMT10A-corrected iPSCs. **A**, Karyostat analysis of isogenic control (1023A, CT) and TRMT10A KO iPSCs showing no chromosomal aberrations in the KO line vs control. B, iCS-digital™ PSC 28-probe test confirming the absence of most frequent chromosomal aberrations in TRMT10A mutant and corrected lines (corrected clone 1 and clone 2). C, Representative immunostaining for OCT4, SSEA4, Nanog and TRA1-60 in TRMT10A KO and corrected iPSCs lines confirming pluripotency. D, Embryoid bodies from TRMT10A KO and corrected lines differentiated into the three germ layers as indicated by SOX17 (endoderm), vimentin (mesoderm) and β-tubulin III (ectoderm) positivity. Scale bar, 40 μm.

**Supplementary Fig. 2.**
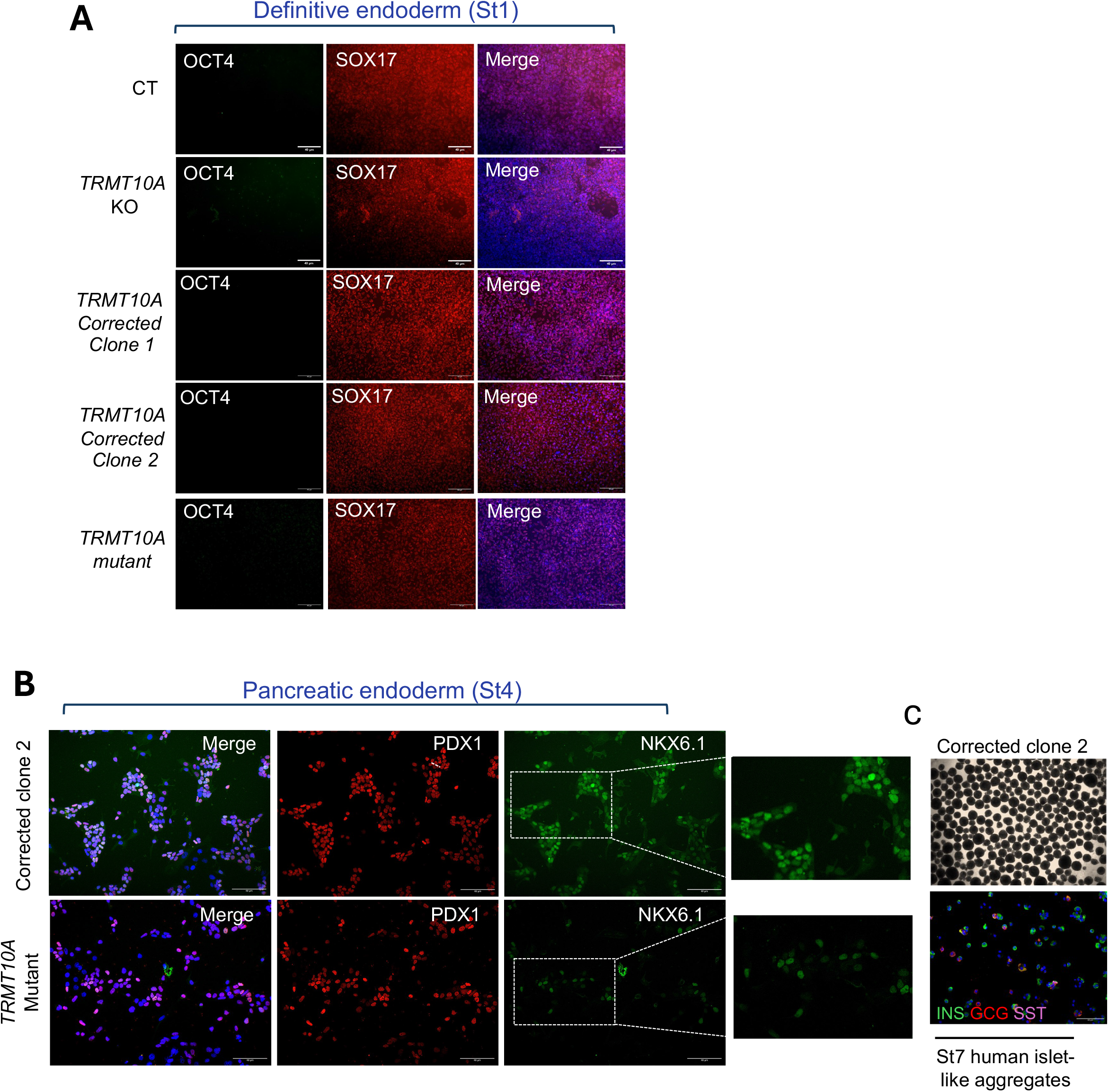
Definitive endoderm and pancreatic endoderm immunostaining in control and TRMT10A-deficient cells. **A**, Representative images of definitive endoderm (St1) immunostaining in isogenic control (CT), *TRMT10A* KO, *TRMT10A* corrected (clone 1 and 2) and *TRMT10A* mutant cells. At St1 the cells were negative for the pluripotency marker OCT4 and positive for the transcription factor SOX17. **B**, Representative immunofluorescence of pancreatic endoderm (St4) in *TRMT10A* corrected (clone 2) and *TRMT10A* mutant cells. Both lines expressed the pancreatic endoderm makers PDX1 and NKX6.1. The dotted lines indicate the portion of the image that is shown magnified on the right side. Scale bar, 40 μm. **c**, Upper panel: bright field representative image of whole aggregates at the end of the differentiation (St7) of *TRMT10A* corrected cells (clone 2). Lower panel insulin (INS), glucagon (GCG) and somatostatin (SST) immunostaining in dispersed aggregates of the same line.

**Supplementary Fig. 3.**
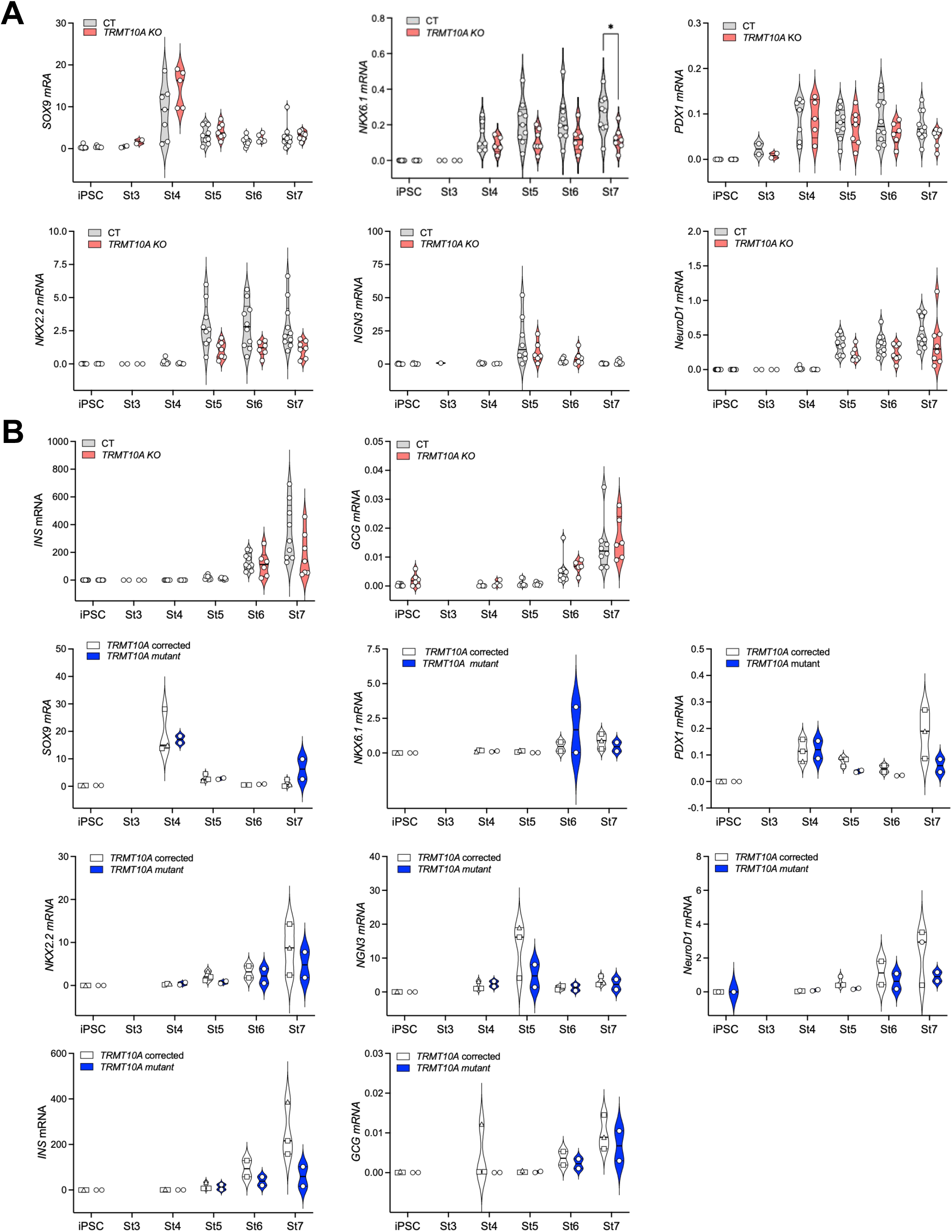
Gene expression profile during iPSC differentiation into human islet-like aggregates. mRNA levels of the differentiation markers *SOX9*, *NKX6.1, PDX1, NKX2.2,* neurogenin3 (*NGN3*), *NeuroD1,* glucagon (*GCG*) and insulin (*INS*) were measured in iPSCs and along differentiation stages 3 to 7 (St3 to St7) in isogenic control (CT) and *TRMT10A KO* **(A)** *or TRMT10A corrected and TRMT10A mutant* **(B)** cells. Data were normalized to the geometric mean of reference genes *ACTB* and *VAPA*. Each point represents an independent separate differentiation. *TRMT10A* corrected clone 1 is represented with squares and *TRMT10A* corrected clone 2 with a triangle. *p<0.05, by Two-way ANOVA with Sidák correction for multiple comparisons.

**Supplementary Fig. 4.**
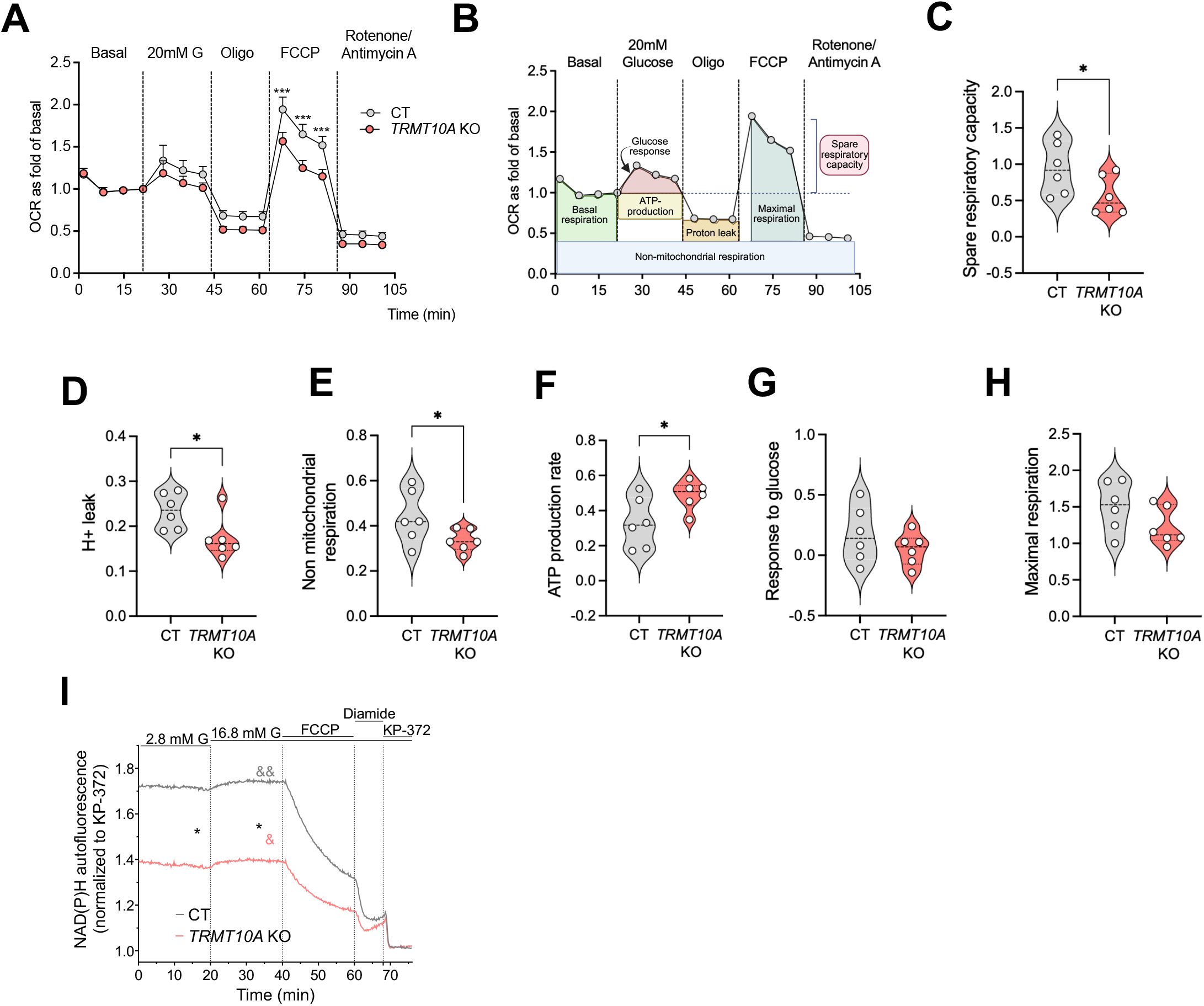
TRMT10A deficiency impairs mitochondrial function. **A,** Mitochondrial function in dispersed St7 islet-like aggregates by Seahorse. The plot shows oxygen consumption rate (OCR) vs time in basal condition (0 mM glucose) and after addition of 20 mM glucose (20 mM G), 5 μM oligomycin (Oligo), 4 μM the uncoupler FCCP, and the mitochondrial poisons rotenone and antimycin (1 μM each). The results are expressed as fold change of the last basal reading. **B**, Schematic representation of the Seahorse assay illustrating key metabolic parameters. **C-H**, Metabolic parameters calculated from the data shown in **A**. **I**, Changes in NAD(P)H autofluorescence during perifusion with 2.8 then 16.8 mM glucose followed by addition of 10 µM FCCP, 200 *s*µM diamide and 10 µM Kp-372 for maximal oxidation of NADH and NADPH. The autofluorescence levels were normalized to the minimal reading measured after addition of Kp-372. Traces show mean +/- SE for 7-9 aggregates from 3-5 independent differentiations. * p<0.05, ***p<0.001, *TRMT10A* KO vs CT, && p<0.01, &&& p<0.001 indicated conditions vs 2.8 mM glucose by two-way ANOVA, followed by Šidák’s correction for multiple comparisons (a,i), or by unpaired t-test (c-h).

**Supplementary Fig. 5.**
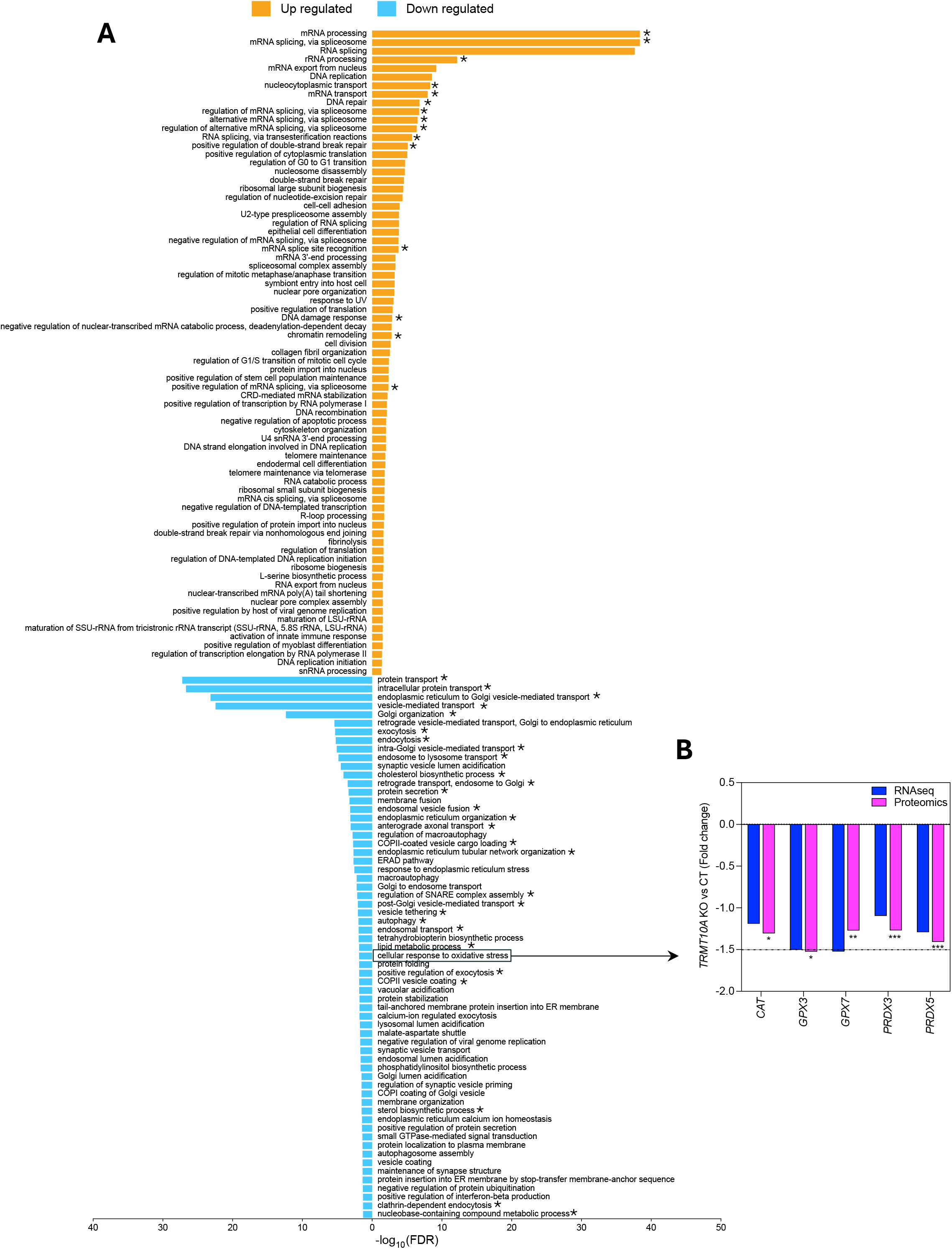
Biological processes dysregulated in TRMT10A KO cells. **A,** DAVID functional gene ontology (GO) analysis performed on the overlapping gene and protein list from RedRibbon. The results highlight upregulated (orange) and downregulated (blue) biological processes in *TRMT10A* KO *St7* aggregates compared to CT***. \**** indicate pathways that were dysregulated in the same direction in *TRMT10A*-silenced EndoC-βH1 cells (Supplementary Fig. 7). **B**, Differentially expressed genes related with cellular responses to oxidative stress.

**Supplementary Fig. 6.**
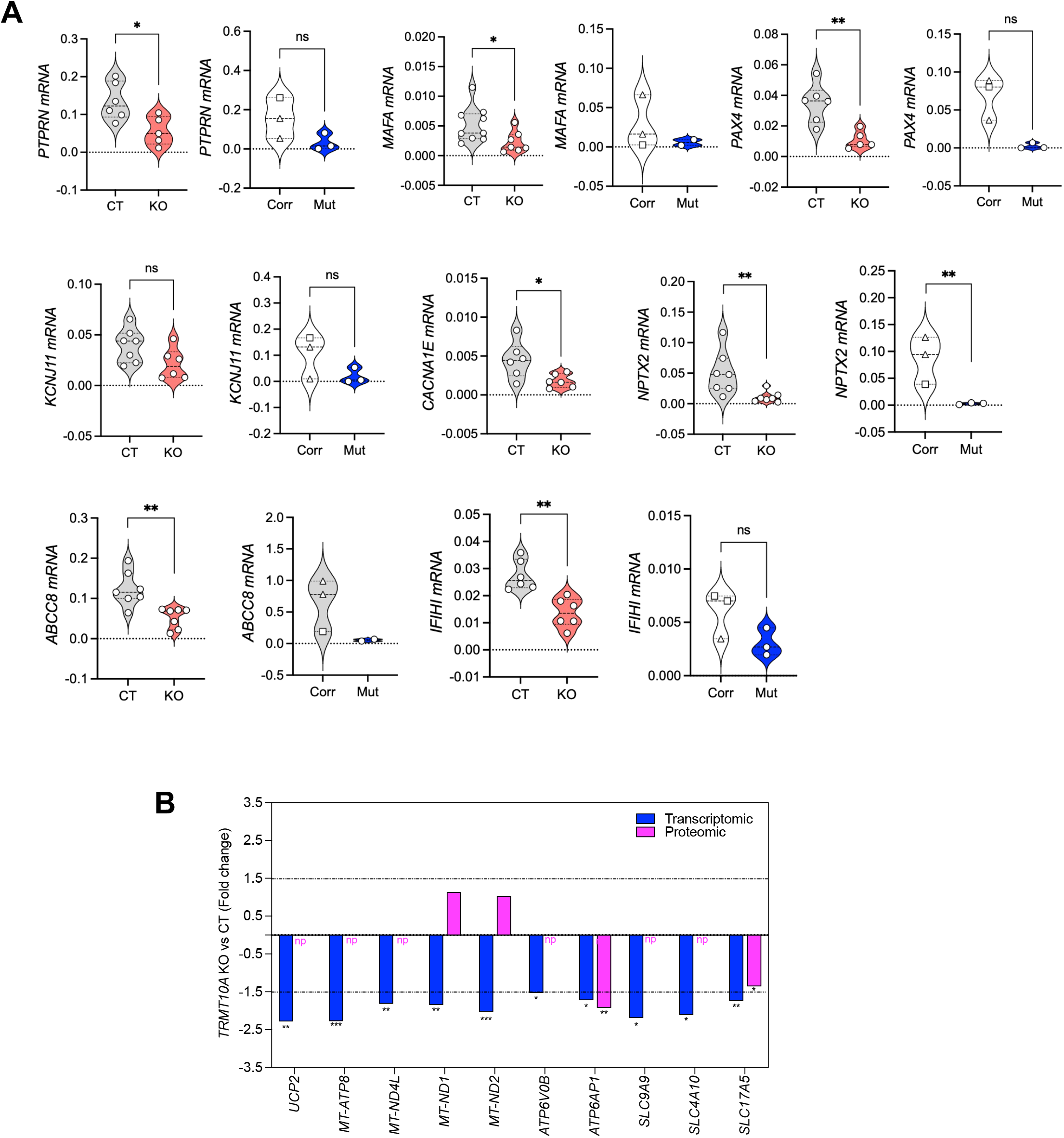
Diabetes, epilepsy and proton transmembrane transport-associated gene expression in TRMT10A-deficient cells. **A**, RT-qPCR validation of diabetes- and epilepsy-associated genes found to be downregulated by RNA sequencing in *TRMT10A*-KO St7 islet-like aggregates. qPCRs done in isogenic control (CT), *TRMT10A* KO (KO), *TRMT10A*-mutant (Mut) and *TRMT10A* corrected (Corr) St7 islet-like aggregates. Data were normalized to *ACTB* and expressed as 2^−ΔCt^. Symbols represent independent differentiations. *TRMT10A* corrected clone 1 is represented with squares and *TRMT10A* corrected clone 2 with a triangle. *p< 0.05, **p<0.01, ns = not significant by unpaired t-test. **B**, Proton transmembrane transport-associated genes identified by GO pathway analysis. Transcriptomic (blue) and proteomic (pink) profiles from the RNA sequencing and proteomic analysis. Differential expression is shown as fold change. Statistical significance was defined as FDR or adjusted p-value < 0.05. np=not present in the proteomic dataset.

**Supplementary Fig. 7.**
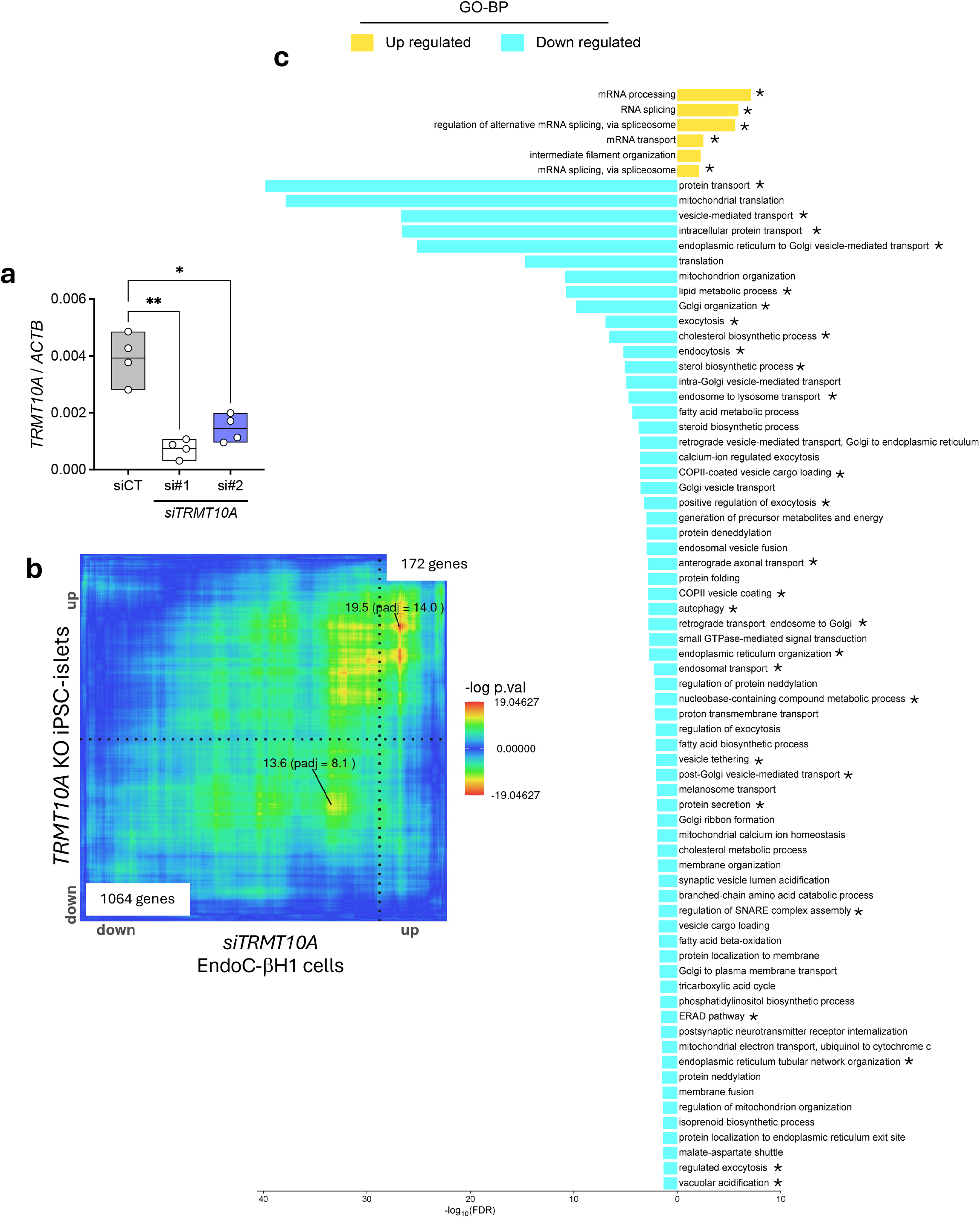
RedRibbon comparison of proteomes of TRMT10A-silenced human β-cells and TRMT10A KO St7 islet-like aggregates. **a,** *TRMT10A* mRNA expression in EndoC-βH1 cells transfected with a control siRNA (siCT) or two siRNAs targeting human *TRMT10A* (si#1 and si#2) n=4 independent experiments used to perform a proteomic analysis. **b,** Rank–rank hypergeometric overlap map generated using RedRibbon. The RedRibbon heatmap indicates the significance (−log10 p-value) of overlap between proteome datasets of *TRMT10A* silenced β-cells (mean of two siRNAs) and *TRMT10A* KO St7 islets. Quadrants represent genes and proteins changing in the same direction (up–up; down–down) or in opposite directions (up–down; down–up) **c**, DAVID functional gene ontology (GO) analysis performed with the up-up (orange) and down-down (light blue) gene sets from the RedRibbon analysis showing the significantly dysregulated biological processes. * indicate pathways that were dysregulated in the same direction at transcriptional and translational level in *TRMT10A* KO St7 islets (Supplementary Fig. 5)

**Supplementary Fig. 8.**
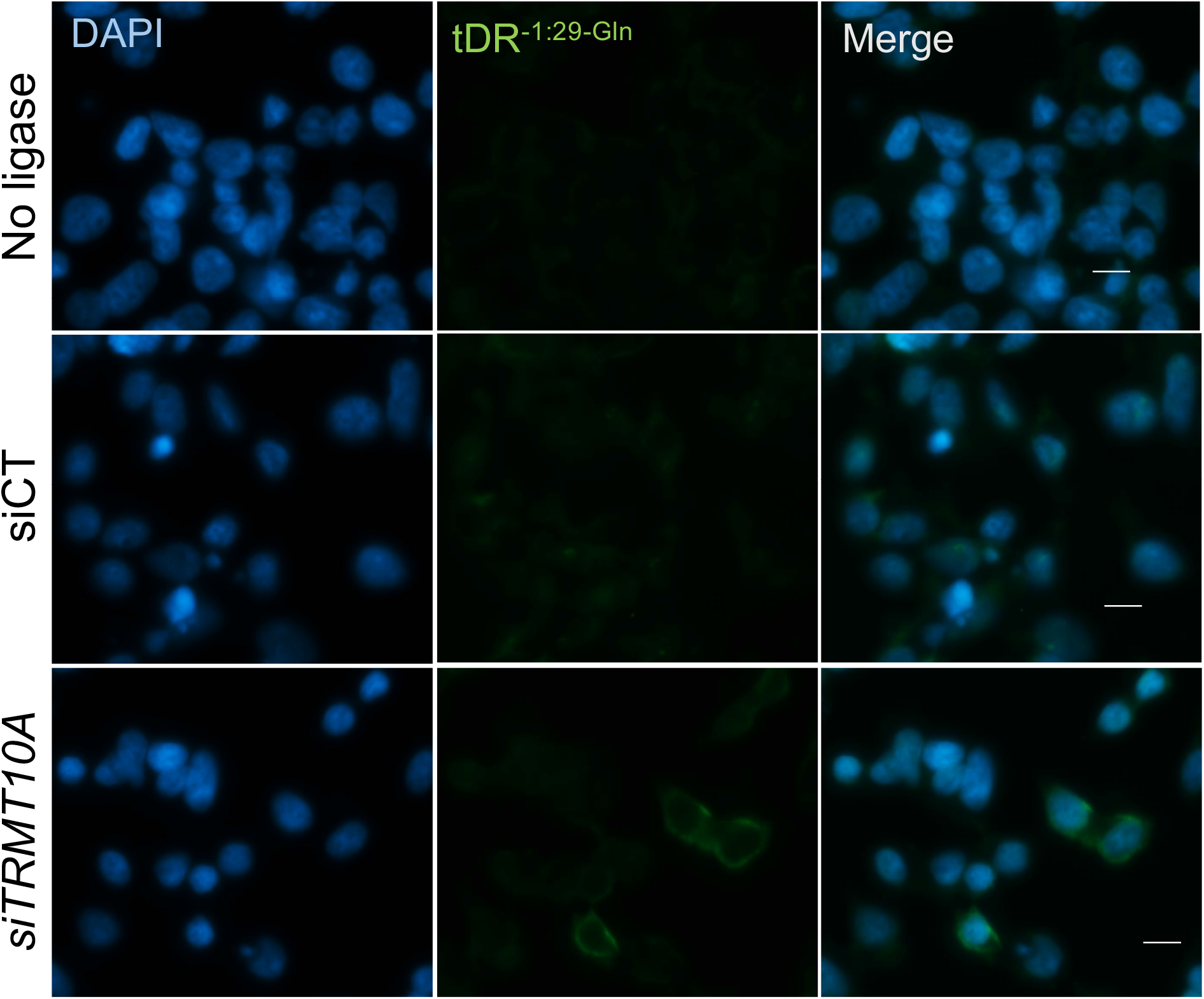
Detection of tDR^-1:29-GlnCTG^ in TRMT10A-silenced EndoC-βH1 cells. Representative LI-FISH images showing tDR^-1:29-GlnCTG^ (green) and nuclei (blue) in EndoC-βH1 cells transfected either with control siRNA (siCT) or siRNA targeting *TRMT10A* (*siTRMT10A*). The “no ligase” condition serves as negative control. Images were acquired using identical exposure settings. Scale bar, 10 µm.

**Supplementary Fig. 9.**
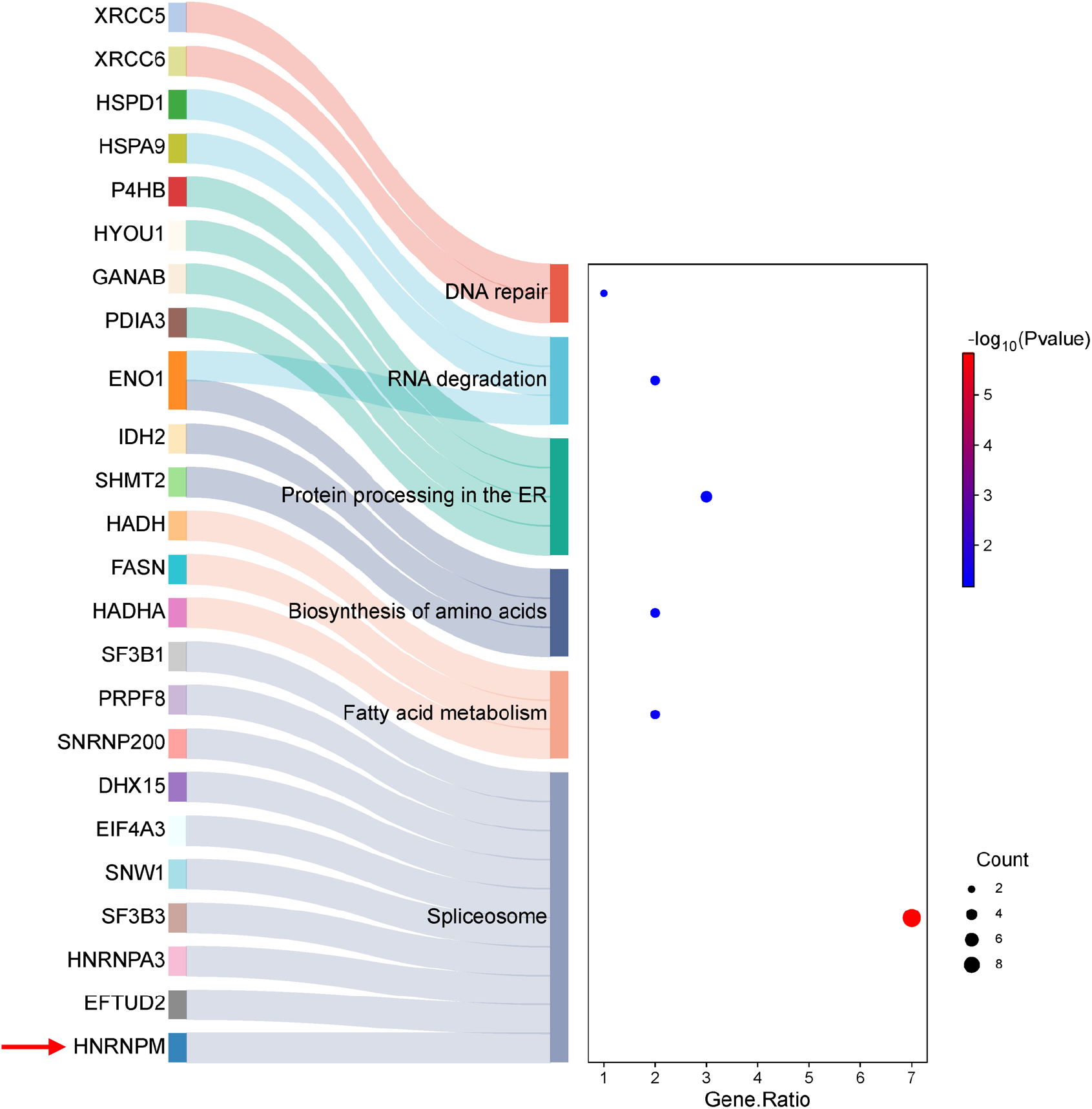
Sankey and dot plot illustrating the proteins potentially interacting with tDR^-1:29-Gln^ in EndoC-βH1 cells, categorized by KEGG pathway analysis

**Supplementary Fig. 10.**
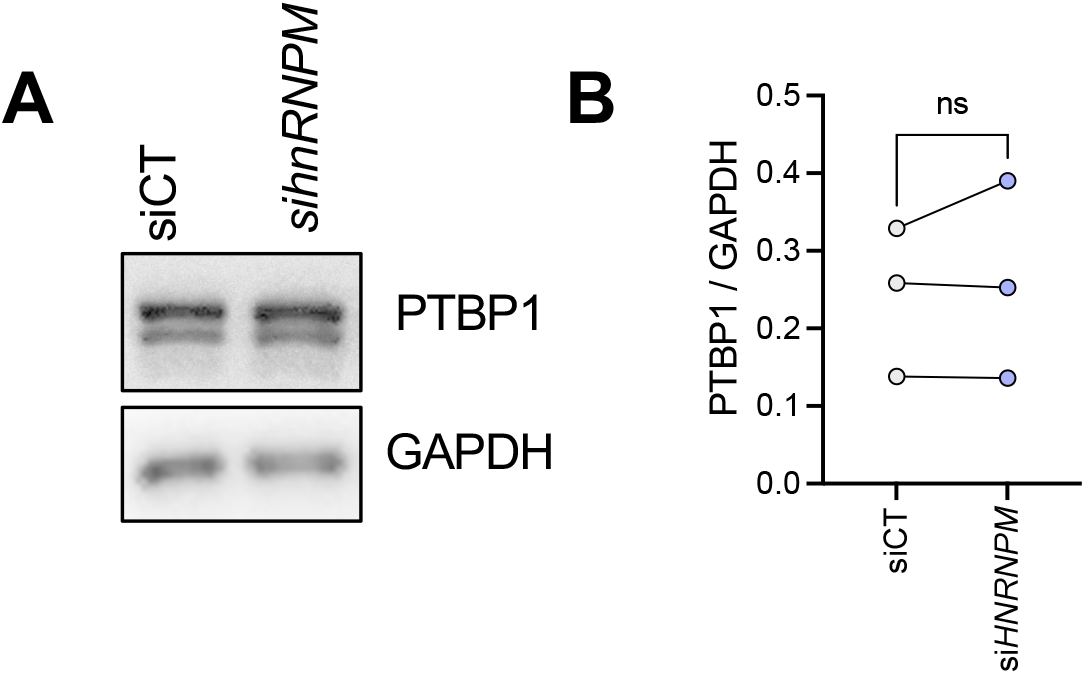
PTBP1 protein expression is not altered in hnRNPM-silencedβ-cells. hnRNPM was silenced in EndoC-βH1 cells and PTBP1 protein expression analyzed by Western Blot. **A,** Representative blot of 3 independent experiments. **B**, Quantification of the blots. ns= not significant

## Supplementary Methods

### iPSCs CRISPR-Cas9-mediated gene editing

The *TRMT10A* KO iPSC cell line was generated by CRISPR-Cas9-mediated-gene editing of the previously validated control iPSC line 1023A(*1*). Correction of the *TRMT10A* mutation (c.379G>A; p.Arg127Stop) was performed in our previously validated iPSC line (HEL122.2, henceforth called *TRMT10A* mutant) from a *TRMT10A*-deficient individual(*2*). Two guide RNAs (gRNA #1 and gRNA #2 for *TRMT10A* KO, gRNA #2 for *TRMT10A* mutant correction) targeting respectively exon 3 and 4 of the *TRMT10A* gene were used. To correct the *TRMT10A* mutation in *TRMT10A-*mutant iPSCs, we introduced a 203 bp donor oligonucleotide (homology-directed repair (HDR) template) encompassing the entire exon 4 and adjacent intronic regions. This donor was designed to (i) correct the c.379G>A mutation, (ii) mutate the PAM site to prevent repeated Cas9 cleavage, and (iii) incorporate a BanII restriction site for clonal selection. Following Cas9-mediated double-strand cleavage upstream of the PAM site, the donor sequence is integrated via HDR, achieving precise mutation correction and enabling downstream selection. The gRNAs and donor sequence were designed and synthesized at Integrated DNA Technologies (IDT), sequences are provided in Supplementary Table 1.

For gene editing iPSCs were plated in Matrigel-coated plates (Corning, #354277) and cultured in Essential 8 medium (Gibco, #A1517001) until reaching 90% confluency. Before nucleofection, a ribonucleoprotein (RNP) complex was assembled by incubating 0.22 nmol of each gRNA with 30 μg Alt-R S.p. HiFi Cas9 Nuclease V3 (IDT, #1081061) and 2μl of R buffer (Neon Transfection System Kit, ThermoFisher, #MPK10096) for 20 minutes at room temperature (RT). iPSC colonies were dissociated into single cells using Accutase (Sigma-Aldrich, #A6964) for 4 min at 37°C, and the reaction was quenched with PBS. Two million cells were then centrifuged at 200 g for 3 min, washed once with PBS and resuspended in 90μl R buffer. The RNP complex, 2μl of Electroporation enhancer (IDT, #227002561), and 4 μg of HDR template (the latter being used only in case of *TRMT10A* correction) were added to the cells, and the mixture was electroporated in a Neon electroporator system (Invitrogen, #MPK5000) with two pulses of 1100 mV 20 ms each. Cells were immediately plated at low density on Matrigel-coated plates containing Stem Flex medium (Gibco, #A3349401) supplemented with 10 μM Rock inhibitor (StemCell, #Y- 27632). For *TRMT10A* correction, 1.7 μl of Alt-R HDR Enhancer V2 (IDT, # 1081061 IDT) were added per ml of medium to improve incorporation of the HDR template. Emerging colonies were hand-picked and expanded. Genomic DNA was extracted using QIAamp DNA Mini Kit (Qiagen, #51304), and the *on-target* regions of gRNA #1 and #2 were amplified by PCR using iProof High-Fidelity DNA Polymerase (BIO-Rad, #172-5301) and sequenced by Sanger sequencing. Following this procedure, one homozygous *TRMT10A* KO clone was identified. For *TRMT10A* mutation correction, positive clones carrying the corrected *TRMT10A* sequence were first identified by restriction enzyme analysis and subsequently confirmed by Sanger sequencing of the on-target region. Two successfully corrected clones were selected for further quality control assessments.

Potential off-target sites predicted by the IDT CRISPR-Cas9 gRNA checker (https://eu.idtdna.com/site/order/designtool/index/CRISPR_SEQUENCE) were analyzed (8 for gRNA#1 and 3 for gRNA#2, Supplementary Table 2). These regions were PCR-amplified and sequenced in the *TRMT10A*-edited clones and their parental iPSC lines. Primer sequences for on-target and off-target analysis are listed in Supplementary Table 3. Chromosomal integrity of *TRMT10A* KO iPSCs was confirmed by KaryoStat Assay (ThermoFisher, #905403). *TRMT10A* corrected clones were assessed for chromosomal integrity by chromosomal microarray analysis (Life&Brain) and/or the iCS-digitalTM PSC 28-probe assay test for detection of the most recurrent chromosomal aberrations in hPSCs (StemGenomics). Pluripotency, and tri-lineage differentiation capacity were assessed by pluripotency marker expression and embryoid body formation as previously described(*2, 3*). Antibodies used are listed in Supplementary Table 4.

### Cell culture

EndoC-βH1 cells(*4*) were cultured in complete EndoC-βH1 culture medium (Supplementary Table 5) on extracellular matrix (ECM, Sigma-Aldrich, #E1270) + fibronectin (Sigma-Aldrich, #F1141)-coated plates at the cell densities specified in Supplementary Table 6.

iPSCs were differentiated into human islet-like aggregates using a previously validated 7-stage protocol(*5*). Islet-like aggregates obtained at St7 were maintained in suspension in Ultra-Low attachment plates (StemCell Technologies, #38071) containing complete St7 medium (Supplementary Table 7) under continuous rotation (100 rpm) in a 5% CO2 incubator. To enhance β-like cell functionality, St7 islet-like aggregates were cultured for one additional month under the same conditions in long-term culture medium (Supplementary Table 8).

Differentiation efficiency was assessed by analyzing the expression of key differentiation markers by immunofluorescence and qPCR, as previously described(*2, 6, 7*). The antibodies and primers used are listed in Supplementary Tables 4 and 9, respectively.

### Dispersion of iPSC-derived islet-like aggregates

St7 or long-term iPSC-derived islet-like aggregates were collected, washed twice with 0.5 mM EDTA, incubated for 8-10 min at 37°C in Accumax^TM^ solution (Sigma-Aldrich, #A7089). Aggregates were gently dispersed by pipetting. Following dispersion, cells were plated on Matrigel-coated plates in either St7 or LTC, as suitable, supplemented with 1 μl/ml Rock inhibitor (StemCell, #Y-27632) at the cell densities described in Supplementary Table 6. After a 16h recovery period, the medium was replaced with complete human islet medium (Supplementary Table 10), and cells were cultured for an additional 24h before any treatment.

### RNA interference

EndoC-βH1 cells were transfected overnight with 30 nM of AllStars Negative Control siRNA (siCT, Qiagen, #1027281), which does not interfere with β-cell function or gene expression(*8*), one of our previously validated siRNAs targeting human TRMT10A(*2, 9*) (siTRMT10A#2), or an siRNA targeting human *hnRNPM* (si*hnRNPM*) using Lipofectamine RNAiMax (Invitrogen, #13778075). Sixteen hours post-transfection, medium was replaced to complete EndoC-βH1 culture medium and cells were processed 96h post-transfection.

Transfection medium composition, lipofectamine concentrations, and siRNA sequences are provided in Supplementary Tables 5, 6 and 11, respectively.

### Transfection with tDR^-1:29-Gln^ mimics

EndoC-βH1 cells were transfected overnight with 60 nM of scramble RNA or tDR^-1:29-Gln^ mimics (Eurogentec, #BA-DN001-001) using Lipofectamine 2000 (Invitrogen, #11668030). Lipofectamine concentration and RNA mimic sequences are provided in Supplementary tables 6 and 12, respectively. Sixteen hours after transfection, medium was changed to complete EndoC-βH1 culture medium, and cells were collected 48h post-transfection.

### RNA extraction, reverse transcription and quantitative PCR

Total RNA was extracted using Direct-zol RNA Miniprep kit (Zymo Research, #R2052) or Direct-zol RNA Microprep Kits (Zymo Research #R2062), as suitable, according to the manufacturer’s instructions. Reverse transcription was performed using 400 ng of total RNA and the Reverse Transcriptase Core Kit (Eurogentec, #RT-RTCK-03) containing random nonamers. RT qPCR was done in a MyiQ2 Two-color Real Time PCR detection system (BIORAD, #170-9790) using SsoAdvanced Universal SYBR® Green Supermix (BIORAD, #1725274). The qPCR results were analyzed using the ΔCt method. *VAPA and/or ACTB* were used as housekeeping genes. The primers used are provided in Supplementary Table 9.

### Quantification of tDR^-1:29-Gln^, tDR^-1:18-Gln^ and full tRNA^-Gln-CTG^ by RT-qPCR

Total RNA was extracted as described above. Prior to reverse transcription, 1-3 µg of RNA was demethylated for 2 h at 37⁰C in a buffer containing 45 mM Tris-HCl (pH 8), 0.9 mM α-ketoglutaric acid 2 Na, 1.8 mM ascorbic acid, 67 µM (NH4)_2_Fe(SO4)_2_, 1.4U/uL RNase inhibitor (40U/uL) (NEB, #M0314S), supplemented with 3 µg of recombinant Escherichia coli AlkB demethylase, and its mutant AlkB-D135S(*10*). Plasmids encoding wild type and mutated enzymes were a gift from Tao Pan (pET30a-AlkB Addgene plasmid #79050 and pET30a-AlkB-D135S Addgene plasmid #79051). Following demethylation, RNA was purified using the Direct-zol RNA Microprep Kit. For full tRNA detection, 100ng of demethylated RNA were reverse-transcribed using the RevertAid First Strand cDNA Synthesis kit (Thermo Scientific, #K1621). qPCR was done in a MyiQ2 Two-color Real Time PCR detection system (BIORAD, #170-9790) using SsoAdvanced Universal SYBR® Green Supermix (BIORAD, #1725274). The qPCR results were analyzed using the ΔCt method. U6 was used as housekeeping gene. The primers used are provided in Supplementary Table 9. For tDR detection, 50 ng of demethylated RNA were reverse-transcribed using the miRCURY LNA RT Kit (Qiagen, #339340) according to the manufacturer’s instructions. Amplification and relative quantification of the target tRNA fragments were performed using the miCURY LNA miRNA PCR assay (Qiagen #339306) and Custom PCR Primer Sets as previously described(*2*) in a MyiQ2 Two-color Real Time PCR detection system. Expression of the miRNA hsa-let-7f-5p (Qiagen, #339306) served as reference. The results were analyzed using the ΔCt method(*2*). Sequences of the analyzed tDRs are provided in Supplementary Table 13.

### Ligation- Fluorescent in situ Hybridization (LI-FISH)

EndoC-βH1 cells were cultured on round 12 mm diameter coverslips (Marienfield, #0111520) and transfected with siRNAs as described above. Dispersed iPSC-islet-like aggregates were cultured on Fluorodish Sterile Culture Dishes, 10 mm well (World Precision Instruments, # FD3510-100). Cell densities and lipofectamine concentrations are described in Supplementary Table 6. At the end of the experiment the cells were fixed with 4% paraformaldehyde (PFA; prepared in PBS) for 45 min on ice and stored in PBS until processing. Permeabilization was performed with 0.5% Triton X-100 (Sigma-Aldrich, # X-100) for 10 min at room temperature. Before stem loop hybridization, wells were washed for 5 min with 0.1x SSC prewarmed to 65°C to denature RNA. Cells were then incubated for 30 min at 45°C in a dry oven with 20 nM denatured stem-loop diluted in *in situ* hybridization buffer (ENZO, #ENZ-33808), followed by 3 washes (5 min each) with 0.1x SSC prewarmed at 50°C. Stem loop ligation was performed using T4 RNA Ligase 1 (New England Biolabs, #M0204S) at 37°C for 1 h according to manufacturer’s indications. After ligation, cells were washed 3 times with 0.1x SSC prewarmed at 50°C and hybridized for 60 min at 60°C with 20 nM digoxigenin-labelled probe diluted in hybridization buffer.

Post hybridization washes included 1 wash with 2x SSC, 0.1 % Tween 20 (Sigma-Aldrich, #P1379) for 5 min, and 3 washes with 0.1x SSC, all prewarmed at 50°C. Cells were then blocked for 45 min at 37°C in a humid chamber with 3 % BSA, 4x SSC, 0.1 % Tween 20. Mouse anti-digoxigenin primary antibody diluted in 3% BSA was applied overnight at 4°C. After 3 PBS washes (5 min each at RT), cells were incubated for 1h a RT with Alexa-647 anti-mouse-secondary antibody and Hoechst 33342 (Sigma-Aldrich, #14533) diluted 1:500 in PBS, followed by 3 washes with PBS. Cells were mounted using VECTASHIELD(R) antifade mounting medium (Vector Laboratories, #VEC.H-1000) was used.

To confirm the specificity of the tDR detection we used a “no-ligase” condition in which the T4 RNA ligase was no added to the reaction. In this situation, the non-ligated stem loop is eventually removed upon washing, and the residual immunofluorescent signal corresponds to the probe binding to the full tRNA or to the remaining non-removed stem loop. This negative control, processed in parallel under identical conditions, showed very little signal compared to the ligase-containing conditions supporting the specificity of the tDR detection.

Images were acquired using an Inverted Zeiss LSM 780 confocal microscope, or a Zeiss Axio Observer 5 inverted microscope with Colibri 5 light source and processed using Image J software and SVI Huygens Pro for colocalization analysis. Stem loop and probe sequences are provided in Supplementary Table 14.

### Subcellular fractionation

Subcellular fractionation of iPSC-derived St7 *TRMT10A* KO islets was performed using the PARIS kit (ThermoFisher Scientific, #AM1921). Briefly, dispersed cells were washed three times with cold PBS1X and collected by centrifugation at 200 g for 5 minutes. Cells were incubated in fractionation buffer for 10 minutes on ice, and then centrifuged at 500 g for 5 minutes at 4°C. The supernatant (cytoplasmic fraction) was collected, and the nuclei pellet was carefully washed twice with fractionation buffer and resuspended in cell disruption buffer.

RNA was demethylated and the localization of tDR^-1:29-Gln^ and tDR^-1:18-Gln^ was assessed by RT-qPCR as described above. 5S rRNA (Qiagen, #339306, YP00203906) and U6 snRNA (Qiagen, #339306, YP02119464) were used as cytoplasmic and nuclear controls, respectively.

### Western Blot

Dispersed iPSC-derived islet-like aggregates or EndoC-βH1 cells were washed twice with ice-cold PBS, and lysed directly in Laemmli buffer 1x. Samples were resolved on 10% SDS-polyacrylamide gels and transferred to methanol-activated PVDF membranes. Membranes were blocked for 1h in 5% non-fat milk prepared in TBS containing 0.1% Tween-20 and then incubated overnight at 4°C with primary antibodies diluted in blocking solution. After 3 washes with TBS 0.1% Tween-20, membranes were incubated for 1h with horseradish peroxidase-conjugated secondary antibodies. Proteins were detected using SuperSignal West Pico PLUS chemiluminescent substrate (ThermoFisher Scientific, #34577) and visualized on a FUSION FX7 Chemiluminescence and Fluorescence imaging system (VILVER). Band intensities were quantified using Fiji. The antibodies and dilutions used are listed in Supplementary Table 4.

### Co-immuno precipitation

EndoC-βH1 cells (10 million) were seeded and cultured overnight in either standard EndoC-βH1 medium (5 mM glucose) or high-glucose medium (20 mM glucose). Cells were harvested and lysed in T7 co-immunoprecipitation buffer (20mM HEPES pH 7.9, 150mM NaCl, 5mM EDTA, 1% NP-40, 1mM DTT, 10% Glycerol) supplemented with RNase inhibitor (1 U/mL; NEB, # M0314S), protease inhibitors, and phosphatase inhibitors. Lysates were precleared for 1h at 4°C using Dynabeads (Invitrogen, #10003D) to minimize nonspecific binding. Following preclearance, lysates were incubated with 2 μg of anti-PTBP1 antibody (Invitrogen, #32-4800) or anti-mouse IgG control antibody (EMD Millipore, #12-371) for 2 h at 25°C. 20 μl of Dynabeads were then added and incubated for additional 30 min at 25°C. Beads were then washed twice with T7 co-IP buffer, twice with low-salt wash buffer (50mM NaCl, 10mM Tris pH 7.5, 0.1% NP-40), and twice with high-salt wash buffer (500mM NaCl, 10mM Tris pH 7.5, 0,1% NP-40). After washing, bound proteins were eluted and analyzed by Western blot.

### Insulin secretion

#### Static non-sequential insulin secretion

Isogenic control and *TRMT10A* KO iPSC-derived St7 islet-like aggregates were incubated for 2h at 37°C in βKREBS without glucose (Human Cell Design, #BK-250) in suspension. Twenty five to thirty aggregates per condition were then transferred to Eppendorf tubes containing 500 μl of ice-cold βKREBS 0.1% BSA supplemented with either 2.8 mM glucose, 16.8 mM glucose, 16.8 mM glucose + 250 μM diazoxide (Sigma-Aldrich, #D9035), 16.8 mM glucose + 25 μM gliclazide (Sigma-Aldrich, #G2167), or 16.8 mM glucose + 30 mM KCl (Sigma-Aldrich, #1.04938). Tubes were kept on ice until all the aggregates were transferred and were then incubated at 37°C for 1h. After this period, the tubes were placed on ice, and the supernatant was collected for insulin secretion measurement.

#### Dynamic Insulin Secretion

Dynamic secretion was assessed using the PERI5LM perifusion instrument (Biorep Technologies). Fifty iPSC-derived islet-like aggregates, maintained in long-term culture for 4 weeks post St7 were used per assay. Aggregates were sequentially perifused at 100μl/min βKREBS + 0.1 % BSAcontaining (i) low glucose (2.8 mM, 16 min), (ii) high glucose (16.8 mM, 24 min), (iii) high glucose + 50 ng/ml Exendin-4 (Sigma-Aldrich, #E7144) (16.6 mM + Ex, 12 min), (iv) low glucose (2.8 mM, 16 min), and (v) low glucose + 30 mM KCl (Merk, #1049380500) (2.8 mM + KCl, 8 min). Samples were collected every 4 minutes through all the experiments.

#### Insulin and proinsulin content

After perifusion or static insulin secretion assay, the aggregates were washed 3 times with ice-cold PBS and lysed in milliQ water by sonication (10 sec ON, 10 sec OFF, 6 cycles, high intensity) using a Bioruptor (CosmoBio). Protein concentration was determined using the Pierce BCA Protein Assay (ThermoFisher Scientific, #23225). For insulin content, 25 μl of cell lysate was mixed with 75 μl of acid ethanol solution (0.18M HCl diluted in 96% ethanol) and incubated ON at 4°C prior to performing the ELISA essay.

Insulin secretion and content were quantified by human insulin ELISA (Mercodia, #10-1113-10), and proinsulin was measured using a human proinsulin ELISA (Mercodia, #10-1118-01) according to manufacturer’s indications.

### NAD(P)H measurements in iPSC-derived islet-like aggregates

For NAD(P)H measurements, iPSC St7 islet-like aggregates were placed in a 37°C temperature-controlled chamber on the stage of an inverted microscope, perifused with Krebs buffer at a flow rate of ∼0.5ml/min, and the autofluorescence of NAD(P)H (λ_ex_ 360 nm; λ_em_ 470 nm) was recorded every 10 sec (*11*). At the end of NAD(P)H autofluorescence recording, the aggregates were exposed to oxidative phosphorylation uncoupler carbonyl cyanide-p-trifluoromethoxyphenylhydrazone (FFCP, 10 µM), thiol oxidizer diamide (200 µM) and NQO1 redox cycling agent 10H-indeno[2,1-e]tetrazolo[1,5-b][1,2,4]triazin-10-one compound with 6H-indeno[1,2-e]tetrazolo[1,5-b][1,2,4]triazin-6-one (1:1) (Kp-372, 10 µM).

### Seahorse assay

Mitochondrial function was assessed by measuring oxygen consumption rate (OCR) in a Seahorse XFp Extracellular Flux Analyzer (Agilent). Dispersed iPSC St7 islet-like aggregates were plated on Seahorse XFp Cell Culture Miniplates (Agilent, #103025-100) at the cell density described in Supplementary Table 6. On the day of the assay, cells were incubated for 1h at 37°C in a non-CO_2_ incubator in KREBS solution(*12*) (20 mM Hepes, 2.5 mM CaCl_2_, 1.16 mM MgS0_4_, 1.2 mM KH_2_PO_4_, 4.7 mM KCl and 114 mM NaCl; pH=7,4) with 0.2% FFA-free BSA. Mitochondrial respiration was measured basally and after sequential injection of KREBS supplemented with 20 mM glucose (final concentration), the ATP synthase inhibitor oligomycin (5 μM), FCCP (4 μM) and electron transport chain inhibitors rotenone and antimycin (1 μM). The last three solutions are part of the Seahorse XFp Cell Mito Stress Test Kit (Agilent, #103010-100). OCR data was normalized to the last basal reading in each sample, considered as 1, and used to calculate different parameters associated with mitochondrial function as follows: *Non mitochondrial oxygen consumption*= Last OCR measured; *Maximal respiration*=First OCR measured after FCCP injection – non mitochondrial oxygen consumption; *H+ (Proton) leak*= Last OCR measured after Oligomycin injection – non mitochondrial oxygen consumption; *ATP production rate*= last OCR of basal measurement – Last OCR after Oligomycin injection; *Spare respiratory capacity*= Maximal respiration – basal respiration; *Acute response*=Last OCR after glucose injection – last OCR of basal measurement.

### Immunofluorescence

EndoC-βH1 cells or dispersed iPSC-derived islet-like aggregates were plated in 12 well removable chamber plates (Ibidi, #81201) at the cell density mentioned in Supplementary Table 6. Cells were fixed with 4% paraformaldehyde for 45 min on ice and stored in PBS until processing. Permeabilization was performed by incubation with 0.5% Triton X-100 (Sigma-Aldrich, # X-100) for 15 min at RT. Before primary antibody incubation, coverslips were blocked with UltraCruz® Blocking Reagent (Santa Cruz Biotechnology, #sc-516214). Cells were incubated ON at 4°C with primary antibodies diluted in PBS containing 0.1%Tween, washed 3 times with PBS at RT, and then incubated for 1h with the corresponding fluorescent secondary antibodies and Hoechst 33342 (Sigma-Aldrich, #14533) diluted 1:500. After 3 washes with PBS, samples were mounted using VECTASHIELD(R) Antifade Mounting Medium (Vector Laboratories, #VEC.H-1000). Primary and secondary antibodies, along with their dilutions, are listed in Supplementary Table 4.

### RNA pull-down followed by mass spectrometry

Potential protein partners of tDR^-1:29-Gln^ were identified using an exploratory RNA pull down assay followed by mass spectrometry, as previously described(*13*). Briefly, 10 million EndoC-βH1 cells were lysed in buffer containing 50 mM Tris–HCl (pH 7.5), 150 mM NaCl, 1% NP40, 0.1% SDS, 0.5% sodium deoxycholate, 1 mM DTT, cOmplete EDTA-free protease inhibitor Cocktail (Roche, #04693132001), and 80 U/ml RNase inhibitor (NEB, #M0314S). Lysis was performed by gentle vortexing followed by 15 min incubation on ice. Lysates were centrifuged at 14000 rpm for 10 min at 4°C, and total protein concentration was determined as previously described. For pull down, 3 μg of either a 5’-biotinylated scramble RNA, or a 5’-biotinylated oligonucleotide mimicking the tDR^-1:29-Gln^ were denatured at 95 °C for 2 min, cooled on ice for 2 min, and refolded at RT for 20 min in 100 μl of RNA structure buffer (10 mM Tris pH 7.0, 0.1 M KCl, 10 mM MgCl_2_). Refolded biotinylated oligonucleotides were then incubated for 1h at RT with continuous rotation with EndoC-βH1 lysate containing 1 mg of total protein. Next, 50 μl prewashed streptavidin-coated magnetic beads, (Dynabeads M-280, Thermo Fisher Scientific, #11205D) were added and incubated for 1h at RT with continuous rotation. Beads were then washed three times with 200 μl washing buffer (50 mM Tris-HCl pH 7.5, 150 mM NaCl, 1 % NP40, 0.1 % SDS, 0.5 % sodium deoxycholate and protease inhibitors). Samples were then sent to BruProMS/APFP proteomics Mass spectrometry facility of the Université Libre de Bruxelles for analysis.

### Sample preparation for proteomic MS analysis

Proteins from iPSC-derived St7 aggregates (n=7 isogenic control and 7 *TRMT10A* KO per group) were extracted in 8M Urea-50mM tris-HCl pH 8.5 using an Ultra-Turrax homogenizer. Lysing Matrix D beads (100mg) were added, and samples were disrupted for 10 min in a TissueLyzer II (Qiagen) at 30 Hz. Samples were further sonicated (3 cycles, 5 sec ON, 5 sec, OFF, amplitude 27%) and centrifuged at 16000xg for 10min. Supernatants were transferred into new tubes, and protein content was determined using the paper method.

For digestion, 50µg of proteins from iPSC-derived St7 control and *TRMT10A* KO islet-like aggregates, or all proteins recovered from beads, were reduced with 10mM TCEP (Merck, #646547) for 30 minutes at RT, then alkylated with 55mM CAA (Merck, #22790) for 30 minutes at RT, protected from light. Samples were then diluted 8-fold with 100mM TEAB (Merck, #18597) and digested ON at 37°C with 1µg of trypsin (Promega, # V5117) and 1µg of LysC (Wako, #129-02541). Tryptic peptides were purified using HLB SPE columns (Affinisep, Spin15- HLB.T3.50) and eluted with ACN: H2O (65:35 v/v) containing 0.1 TFA (Trifluoroacetic acid). After evaporation, samples were reconstituted in H_2_O with 0.1 % formic acid (FA) before injection.

### MS acquisition for proteomics analysis

RNA pull-down peptides

Peptides obtained from RNA pull down assays were resuspended in 100% H2O/0.1% FA. Eight microliters of the resuspended peptides (80%) were separated on a ChromXP C18 CL column (150 x 0.3mm, 3µm, Sciex) using a two steps acetonitrile gradient: 5-35% ACN / 0.1% FA over 5min, followed by 35-80% ACN / 0.1% FA over 3min. Peptides were then sprayed online on a Triple TOF 5600 mass spectrometer (Sciex, Concord, Canada) coupled to an EK425 HPLC System (Eksigent, Dublin, CA), and data were acquired using Data-Dependent-Acquisition (DDA) mode. MS1 spectra were acquired from 400-1250 m/z with a 250ms accumulation. The 20 most intense precursors (charge states 2 to 4) were selected for fragmentation, and MS2 spectra were collected from 50-2000 m/z with a 200ms of accumulation time in high-sensitivity mode. Precursor ions were excluded for reselection for 12s.

#### Peptides from iPSC-derived St7 islet-like aggregates

Peptides purified from iPSC-derived ST7 cells were separated on a Kinetex XB-C18 column (150 x 0.3mm, 2.6µm; Phenomenex) using a two steps acetonitrile gradient: 5-25% ACN / 0.1% FA over 65 min, followed by 25-60% ACN / 0.1% FA over 25min, and then sprayed online in a Triple TOF 5600 mass spectrometer (Sciex, Concord, Canada) coupled to an EK425 HPLC system (Eksigent, Dublin, CA) using SWATH-DIA acquisitions. Seventy-one variable isolation windows were used to cover a mass range of 400-1250 m/z. MS2 spectra were collected from 50-2000 m/z. Collision energy for each window was calculated for a 2+ charge ion centered in the window with a spread of 15. Fragment-ions scans were acquired in high-sensitivity mode with 45 ms accumulation time, and survey scans were acquired in high-resolution mode at the start of each cycle, resulting in a duty cycle of ∼3.3s.

### MS data analysis

All raw.Wiff/Wiff.scan data files were converted to mzML file format with MSConvert (Version: 3.0). Converted mzML files from RNA pull down assay were analysed using Fragpipe computational platform (v18.0) with MSfragger (v3.5), (*14, 15*) Philosopher (v4.4.0)(*16*). Peptide identifications were obtained using MSFragger search engine on .mzML files, using a protein sequence database of human (UP000005640) from Uniprot (downloaded 07th April, 2022, containing “sp” and “tr” sequences, no isoforms) supplemented with common contaminant proteins and reversed protein sequences as decoys. Mass tolerances for precursors and fragments were set to 30 and 20 ppm respectively, and with spectrum deisotoping mass calibration 4, and parameter optimization enabled. Enzyme specificity was set to “trypsin” with enzymatic cleavage, and a maximum of 5 missed trypsin cleavages were allowed. Isotope error was set to 0/1/2. Peptide length was set from 6 to 50, and peptide mass was set from 500 to 5000 Da. Variable modifications (methionine oxidation, acetylation of protein N-termini, deamidation of Q and N [+0.984016Da], biotin on lysine [+226.0776], pyro-Glu/Gln [-18.0106/-17.0265 Da], and carbamylation on lysine and protein terminus [+43.005814]) were added while carbamidomethylation of Cysteine was set as a fixed modification. The maximum number of variable modifications per peptide was set to 5.

MS/MS search results were further processed using the Philosopher toolkit with PeptideProphet (with options for accurate mass model binning, semi-parametric modeling with computation of possible non-zero probabilities for decoy entries) and with ProteinProphet. Further filtering to 1% protein-level FDR allowing unique and razor peptides were used, and final generated reports were filtered at each level (PSM, ion, peptide, and protein) at 1% FDR.

Converted mzML files from iPSC samples were analyzed with DIA-NN; (version 1.8.2 beta 27 (*17*) with search parameters set as follows: precursor FDR 1%; mass accuracy at MS1 and MS2 both set to 0; scan window set to 0; isotopologues and MBR turned on; protein inference at gene level; heuristic protein inference enabled; quantification strategy set to QuantUMS (high accuracy); neural network classifier double-pass mode; cross-run normalization RT-dependent. DIA spectra were searched against a prior in house generated library containing 12410 proteins covered by 189947 peptides including common contaminants (*18*) and protein re-annotation was performed

Spectral library was generated in house using DDA and DIA data acquired on tT5600 using a similar separation LC gradient via FragPipe (version 18.0) using a pre-defined workflow DIA_SpecLib_Quant. Specifically, MS Spectra were searched as described above for DDA data with MSFragger (version 3.5), using a database containing human protein (UP000000589, 54739 reviewed and unreviewed entries, downloaded from UniProtKB on 05th October 2023) supplemented of common contaminants and decoys. In the validation step, MSBooster was implemented on both spectra and RT levels, and then Percolator and ProteinProphet integrated in Philosopher (version 4.4.0)(*16*) were used for PSM validation and protein inference. Library generation was conducted using EasyPQP (IonQuant version 1.9.8) with RT calibration based on ciRTs and Lowess fraction set to 0.04. Only fragment types b and y were included with a tolerance of 15 ppm with a max delta_unimod.

The protein group output matrix and associated experiment annotation files were imported into FragPipe-Analyst interface for differential abundance testing(*19*). The minimum percentage of non-missing values was set to 60% in at least one condition. A cutoff of the adjusted value of p < 0.05 (Benjamini-Hochberg method) along with an absolute log2 fold change of 0.58 was applied to determine significantly dysregulated proteins.

#### Proteomic analysis in *TRMT10A*-silenced EndoC-βH1 cells

*TRMT10A* expression was silenced by RNA interference in EndoC-βH1 cells (n=4 independent experiments) using two previously validated *TRMT10A* siRNAs (siTRMT10A#1 and #2)(*2*). Cells transfected with a control siRNA (siCT, Qiagen, #1027281), which does not interfere with β-cell function or gene expression(*8*) were used as control. siRNA sequences are provided in Supplementary Table 11. A total of 600,000 cells per condition (*TRMT10A*-silenced or not) were solubilized in 500 µL lysis buffer (8 M urea, 20 mM HEPES pH 8.0) and lysed by sonication (Microtip, 15 W output, 3 bursts of 15 s each). Lysates were cleared by centrifugation (20,000 x g for 15 min at 21°C). Cysteines were reduced and subsequently alkylated by adding 15 mM dithiothreitol for 30 min at 55°C and 30 mM 2-iodoacetamide for 15 min at room temperature in the dark. Protein concentration was determined by Bradford assay. A total of 150 µg of protein material was isolated and diluted two-fold in 20 mM HEPES, pH 8.0 <u>Protein digestion:</u> The samples were then digested with 2.0 µg Lysyl endopeptidase (endo-Lys C, WAKO) for 90 min at 37°C, after which they were further twofold diluted and digested overnight with 2.0 µg trypsin (Promega).

The peptide mixture was then acidified with trifluoroacetic acid (TFA), centrifuged for 15 min at 1,780 x g, and the supernatant was cleaned in SampliQ Columns, using 60% acetonitrile (ACN), 0.1% TFA as elution solvent. The samples were then completely dried and stored at -20°C until analysis.

<u>SCX fractionation:</u> samples were resuspended in 100 µL 2% ACN, 1% TFA and loaded on a wetted and equilibrated SCX tip (made in-house). SCX is a strong cation exchange sorbent used to extract basic analytes from aqueous samples. Fractionation was done by applying 20-40µL of each fractionation buffer subsequently on the tip and collecting the eluent. A total of four fractions were collected, dried, resuspended in 20 µL loading solvent A (2% ACN, 0.1% TFA) and analyzed by liquid chromatography-mass spectrometry (LC-MS/MS).

- 100 mM ammonium acetate, 20% ACN, 0.5 % formic acid (FA)
- 175 mM ammonium acetate, 20% ACN, 0.5 % FA
- 375 mM ammonium acetate, 20% ACN, 0.5 % FA
- Elution buffer: 80% ACN, 5% ammonium hydroxide

<u>LC-MS/MS:</u> 8 µL of the peptide mixture was analyzed by an Ultimate 3000 RSLC nano LC (Thermo Fisher Scientific, Bremen, Germany) in-line connected to a Q Exactive HF mass spectrometer (Thermo Fisher Scientific). The peptides were first loaded on a trapping column (made in-house, 100 μm internal diameter (I.D.) × 20 mm, 5 μm beads C18 Reprosil-HD, Dr. Maisch, Ammerbuch-Entringen, Germany) and after flushing from the trapping column, peptides were loaded on an analytical column (made in-house, 75 μm I.D. × 400 mm, 1.9 μm beads C18 Reprosil-HD, Dr. Maisch) using a non-linear 150 min gradient of 2-56% solvent B (0.1% formic acid in water/acetonitrile, 20/80 (v/v)) at a flow rate of 250 nL/min. This was followed by a 10 min wash reaching 99% solvent B’ and re-equilibration with solvent A (0.1% FA in water). Column temperature was kept constant at 50°C. The mass spectrometer was operated in data-dependent, positive ionization mode, automatically switching between MS and MS/MS acquisition for the 16 most abundant peaks in a given MS spectrum. The source voltage was set to 3.0 kV and the capillary temperature was 275°C. One MS1 scan (m/z 375-1500, AGC target 3E6 ions, maximum ion injection time of 60 ms) acquired at a resolution of 60,000 (at 200 m/z) was followed by up to 16 tandem MS scans (resolution 15,000 at 200 m/z) of the most intense ions fulfilling predefined selection criteria (AGC target 1E5 ions, maximum ion injection time of 80 ms, isolation window of 1.5 m/z, fixed first mass of 145 m/z, spectrum data type: profile, under fill ratio 2%, intensity threshold 1.3E4, exclusion of unassigned, singly charged precursors, peptide match preferred, exclude isotopes on, dynamic exclusion time of 12 s). The HCD collision energy was set to 30% Normalized Collision Energy.

Data analysis: Database searching was performed with MaxQuant (version 1.5.8.3) using the integrated Andromeda search engine with default search settings including a false discovery rate set at 1% on both the peptide and protein level. Spectra were searched against human proteins in the UniProt/Swiss-Prot database (database release version of November 2016). The mass tolerance for precursors and fragment ions was set to 20 and 4.5 ppm, respectively, during the main search. Enzyme specificity was set to C-terminal to arginine and lysine, also allowing cleavage at arginine/lysine-proline bonds with a maximum of two missed cleavages. Variable modifications were set to oxidation of methionine (to sulfoxides) and acetylation of protein N-termini. A minimum of one peptide in total was required for identification. Downstream data processing was done using the Perseus software (v1.5.8.5). A total of 6,005 protein groups were successfully identified after removing proteins that were only identified with a modified peptide, reversed hits and potential contaminants. Log2 fold changes between *TRMT10A*-silenced and siCT were calculated. Proteins with less than 4 valid values per comparison were removed.

### RNA sequencing

RNA sequencing was performed in isogenic control (n=8 differentiations) and *TRMT10A* KO (n=8 differentiations) St7 iPSC-derived islet-like aggregates. Total RNA was isolated using Direct-zol RNA Miniprep kit (Zymo Research, #R2052), quantified by nanodrop, and assessed for quality using an Agilent Bioanalyzer (Agilent) with the RNA 6000 nano kit (Agilent, #5067-1511). Samples with RNA integrity number (RIN) ≥7.6 were used for sequencing. Library preparation (TruSeq Stranded Total RNA, Illumina) and sequencing were performed by the Lausanne Genomic Technologies Facility, University of Lausanne Switzerland. Data analysis was conducted using the QIAGEN CLC Genomics Workbench (RNA seq Analysis portal 5.1). Trimmed reads were mapped to the human reference genome GRCh38.103. Differential gene expression (DE) was determined using a cutoff of FDR p < 0.05 along with an absolute log2 fold change of 0.58. Pathway enrichment analysis was performed using DAVID (Database for Annotation, Visualization, and Integrated Discovery)(*20, 21*). Graphical representations of the RNA sequencing data were generated using SRplot(*22*).

---

**Supplementary Table 1:**
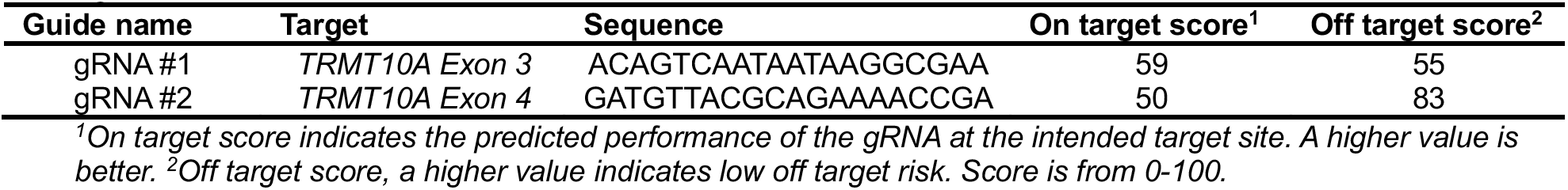
sequence of the gRNAs used for CRISP-Cas9-mediated TRMT10A editing.

**Supplementary Table 2:**
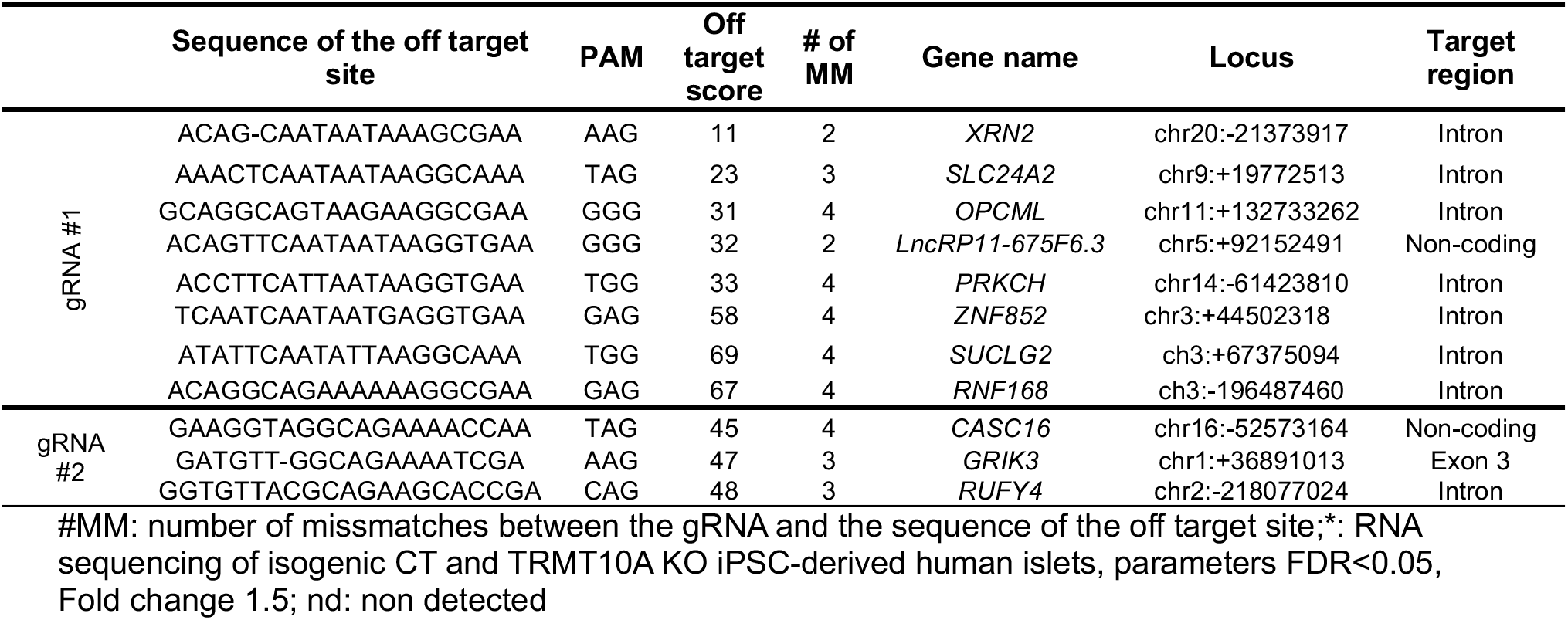
List of potential off targets for gRNA #1 and gRNA #2.

**Supplementary Table 3:**
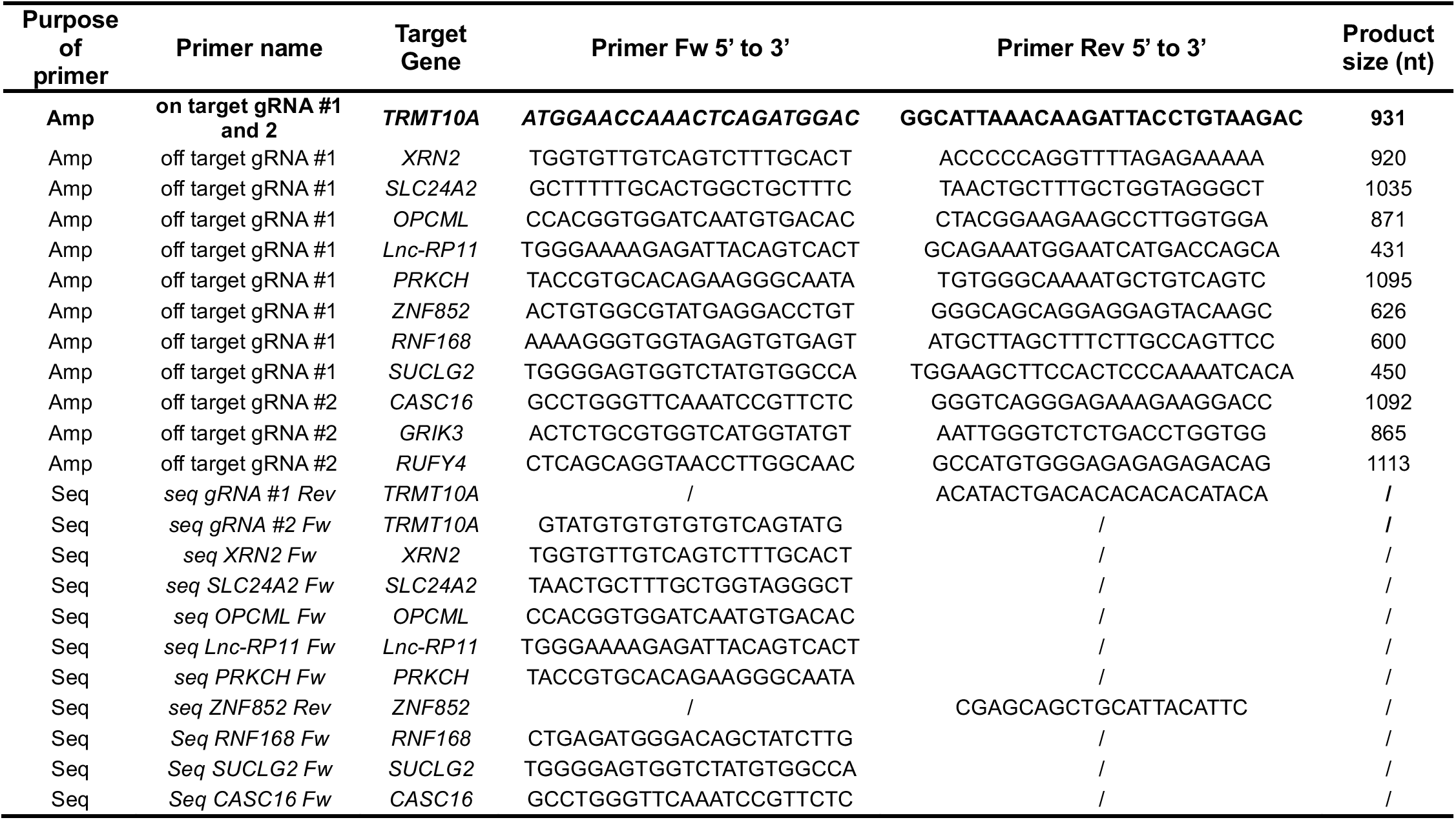
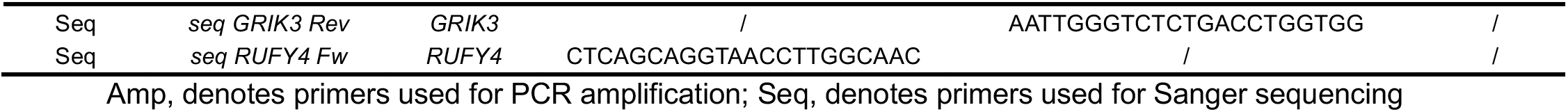
sequence of primers used to amplify and sequence the on target and off target regions of gRNA #1 and gRNA #2.

**Supplementary Table 4:** antibodies used.

| Peptide/protein target | Antibody name | Manufacturer, catalogue # | Species and clonality | Dilution | RRID |
| --- | --- | --- | --- | --- | --- |
| GAPDH | Human Anti-G3PDH antibody | Trevigen, # TRE-2275-PC-020 | Rabbit polyclonal | 1:1000 | N/A |
| α-Tubulin | Anti-TUBA4A (TUBA1) antibody | Sigma-Aldrich, # T9026 | Mouse monoclonal | 1:5000 | AB_477593 |
| TRMT10A | TRMT10A polyclonal antibody | Proteintech, # 17294-1-AP | Rabbit polyclonal | 1:000 | AB_2878377 |
| hnRNPM | HNRNPM Polyclonal antibody | Proteintech, # 26897-1-AP | Rabbit polyclonal | WB: 1:1000<br>IF: 1:200 | AB_2880673 |
| PCSK1 | Anti-PC1/3 antibody | Abcam, # ab220363 | Rabbit monoclonal | 1:1000 |  |
| PTBP1 | PTBP1 Monoclonal Antibody | Invitrogen, #32-4800 | Mouse monoclonal | WB: 1:1000<br>CO-IP: 1:500 | AB_2533082 |
| IgG | Normal mouse IgG | Sigma Aldrich, #12-371 | Mouse polyclonal | CO-IP: 1:500 |  |
| Digoxigenin | anti-Digoxigenin | Roche, #11333062910 | Mouse | 1:250 | AB_2313639 |
| OCT-3/4 | Oct-4A (C30A3) Rabbit mAb | Cell signalling, # 2840 | Rabbit Monoclonal | 1:400 | AB_2167691 |
| SSEA-4 | SSEA4 Monoclonal Antibody (MC-813-70) | ThermoFisher Scientific, # MA1-021 | Mouse Monoclonal | 1:500 | AB_2536687 |
| TRA-1-60 | TRA-1-60 Monoclonal Antibody (TRA-1-60) | ThermoFisher Scientific, # MA1-023 | Mouse Monoclonal | 1:100 | AB_2536699 |
| Nanog | Nanog (D73G4)XP Rabbit mAb | Cell signalling, # 4903 | Rabbit, Mono clonal | 1:400 | AB_10559205 |
| Vimentin | Anti-Vimentin antibody | Abcam, # ab137321 | Rabbit polyclonal | 1:250 | AB_2921312 |
| Beta tubulin III | Anti-β III Tubulin mAb | Promega, # G7121 | Mouse monoclonal | 1:200 | AB_430874 |
| SOX17 | Human SOX17 antibody | R&D systems, # AF1924 | Polyclonal Goat IgG | 1:400 |  |
| Insulin | Polyclonal Guinea Pig Anti-Insulin antibody | DAKO, # A056401-2 | Guinea pig polyclonal | 1:2 | AB_2617169 |
| Glucagon | Anti-Glucagon-Antibody produced in rabbit | Sigma-Aldrich, # SAB4501137 | Rabbit polyclonal | 1:1000 | AB_10761583 |
| Somatostatin | Anti somatostatin antibody | Abcam, # ab108456 | Rabbit polyclonal | 1:1000 | AB_11158517 |
| PDX1 | Human PDX-1/IPF1 Antibody | R&D systems, #AF2419 | Polyclonal Goat IgG | 1:500 |  |
| NKX6.1 | Purified Mouse Anti-Nkx6.1 | BD Bioscience, #563022 | Mouse monoclonal | 1:250 |  |
| Pro-Insulin | Human pro-insulin antibody produced in mouse | DSHB, #GS-9A8 | Mouse monoclonal | 1:200 | AB_532383 |
| Anti-mouse-peroxidase IgG | Peroxidase AffiniPure F(ab')<sub>2</sub> Fragment Donkey Anti-Mouse IgG (H+L) | Jackson ImmunoResearch, # 715-036-150 | Rabbit polyclonal | 1:5000 | AB_2340773 |
| Anti-rabbit-peroxidase IgG | Peroxidase AffiniPure F(ab')<sub>2</sub> Fragment Donkey Anti-Rabbit IgG (H+L) | Jackson ImmunoResearch, # 711-036-152 | Rabbit polyclonal | 1:5000 | AB_2340590 |
| Anti-guinea-pig Alexa Fluor 488 IgG | Alexa Fluor 488 AffiniPure Donkey anti-Guinea Pig IgG (H + L) | Jackson ImmunoResearch, # 706-545-148 | Donkey polyclonal | 1:500 | AB_2340472 |
| Anti-rabbit Alexa Fluor 488 IgG | Alexa Fluor 488 AffiniPure Donkey anti-rabbit IgG (H + L) | Jackson ImmunoResearch, # 711-545-152 | Donkey polyclonal | 1:500 | AB_2313584 |
| Anti-rabbit RRX IgG | Rhodamine Red-X (RRX) AffiniPure Donkey Anti-Rabbit IgG (H+L) | Jackson ImmunoResearch, # 711-295-152 | Donkey polyclonal | 1:500 | AB_2340613 |
| Anti-Mouse Alexa Fluor 488 IgG | Alexa Fluor 488 AffiniPure Donkey Anti-Mouse IgG (H+L) | Jackson ImmunoResearch, # 715-545-151 | Donkey polyclonal | 1:500 | AB_2341099 |
| Anti-Mouse Rhodamine IgG | Rhodamine (TRITC) AffiniPure Donkey Anti-Mouse IgG (H+L) | Jackson ImmunoResearch, # 715-025-151 | Donkey polyclonal | 1:500 | AB_2340767 |
| Anti-Mouse Alexa Fluor 647 IgG | Alexa Fluor 647-AffiniPure Donkey Anti-Mouse IgG (H+L) | Jackson ImmunoResearch, # 715-605-151 | Donkey polyclonal | 1:200 | AB_2340863 |
| Anti-Goat Alexa 488 IgG | Alexa Fluor® 488 AffiniPure™ Donkey Anti-Goat IgG (H+L) | Jackson ImmunoResearch, # 705-545-147 | Donkey polyclonal | 1:500 | AB_2336933 |
| Anti-Mouse IgG Light Chain Specific (HRP) | Rabbit Anti-Mouse IgG (Light Chain Specific) (D3V2A) Monoclonal Antibody (HRP Conjugate) | Cell Signaling, #58802 | Rabbit polyclonal | 1:10000 |  |
| Anti-Rabbit IgG Light Chain Specific (HRP) | Mouse monoclonal [SB62a] Anti-Rabbit IgG light chain (HRP) | Abcam, #ab99697 | Mouse monoclonal | 1:10000 |  |

**Supplementary Table 5:** Mediums for EndoC-βH1.

| EndoC-βH1 culture medium |  |
| --- | --- |
| Component (final concentration) | Brand and catalog number |
| DMEM, low glucose, pyruvate | Gibco, #31885023 |
| 5.5 µg/ml human transferrin | Sigma-Aldrich, # T8158 |
| 6.7ng/ml sodium selenite | Sigma-Aldrich, # S1382 |
| 2 % Bovine Serum Albumin (BSA) Fraction V Free Fatty acid (FFA)-free | Roche, # 10775835001 |
| 10 mM Nicotinamide | Sigma-Aldrich, # N0636 |
| 50 µM beta-2-mercaptoethanol | Sigma-Aldrich, # M3148 |
| 100 µg/ml streptomycin, 100 U/ml penicillin | Gibco, # 15140122 |
| EndoC-βH1 transfection medium |  |
| DMEM, low glucose, pyruvate | Gibco, #31885023 |
| 5.5 µg/ml human transferrin | Sigma-Aldrich, # T8158 |
| 6.7ng/ml sodium selenite | Sigma-Aldrich, # S1382 |
| 10 mM Nicotinamide | Sigma-Aldrich, # N0636 |
| 50 µM beta-2-mercaptoethanol | Sigma-Aldrich, # M3148 |
| Fetal Bovine serum (FBS) 2% | Gibco, # A5256701 |

**Supplementary Table 6:** number of cells seeded per well and volume of transfection reagent.

| Cell type | Type of plate and application | Cell number per well | Volume of lipofectamine RNAiMax per well | Volume of lipofectamine 2000 per well |
| --- | --- | --- | --- | --- |
| EndoC-βH1 cells | 24 well plate | 200,000 | 1 µl in 500 µl | 1 µl in 500 µl |
|  | Round coverslips | 200,000 | 1 µl in 500 µl | / |
| Dispersed iPSC-derived islets | 24 well plate | 250,000 | / | / |
|  | 12 well chamber, removable, plates (Idibi) | 50,000 | / | / |
|  | Fluorodish Sterile Culture Dishes | 50,000 | / | / |
|  | 8 well Seahorse plate | 50,000 | / | / |

**Supplementary Table 7.**

| Composition of Stage 7 medium |  |  |
| --- | --- | --- |
|  | Component (final concentration) | Brand and catalog number |
| Basal 3 | MCDB131, no glutamine | Gibco, # 10372-019 |
|  | 2 mM GlutaMAX | Gibco, # 35050-038 |
|  | 1,5 g/l NaHCO<sub>3</sub> | Merk Millipore, # 1.06329.0500 |
|  | 20 mM Glucose | Sigma-Aldrich, # G8769 |
|  | 0.5 X Insulin-Transferin-Selenium-X (ITS-X 100X) | Gibco, # 51500056 |
|  | 2% BSA Fraction V FFA-Free | Roche, # 10775835001 |
|  | 10 µg/ml Heparin | StemCell Technologies, # 07980 |
|  | 10 µM Zinc Sulfate | Sigma-Aldrich, # Z0251 |
| Supplements for Basal 3 | 1 µM GC1 | Tocris, # 4554 |
|  | 10 µM Trolox | Sigma-Aldrich, # 238813-1G |
|  | 20 µM JNKi | Selleckchem, # SP600125 |
|  | 75 µM Resveratrol | Sigma-Aldrich, # R5010 |
|  | 2 µM R428 | Selleckchem, # S2841 |
|  | 1 mM N-acetyl L-cysteine | Sigma-Aldrich, # A9165 |
|  | 50U/ml Penicillin/Streptomycin | Gibco, # 15140122 |

**Supplementary Table 8.**
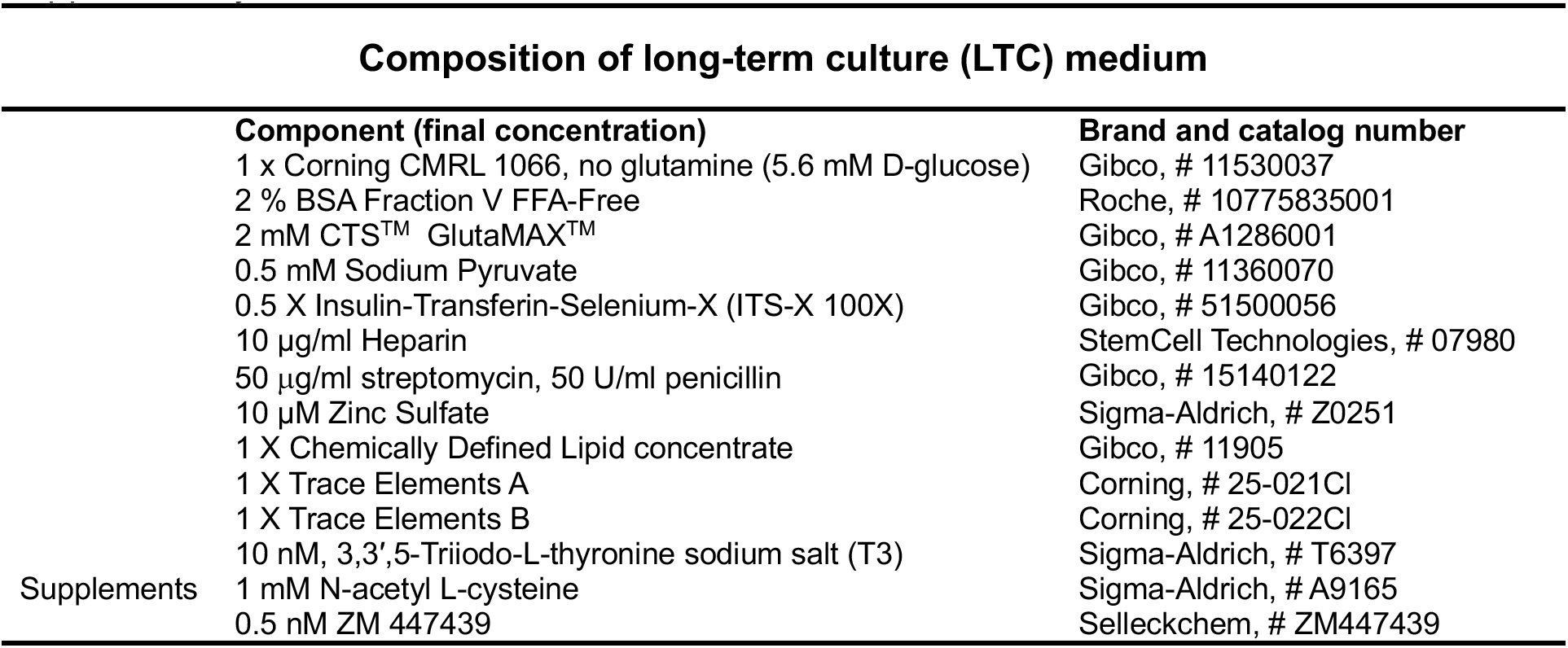

**Supplementary Table 9:**
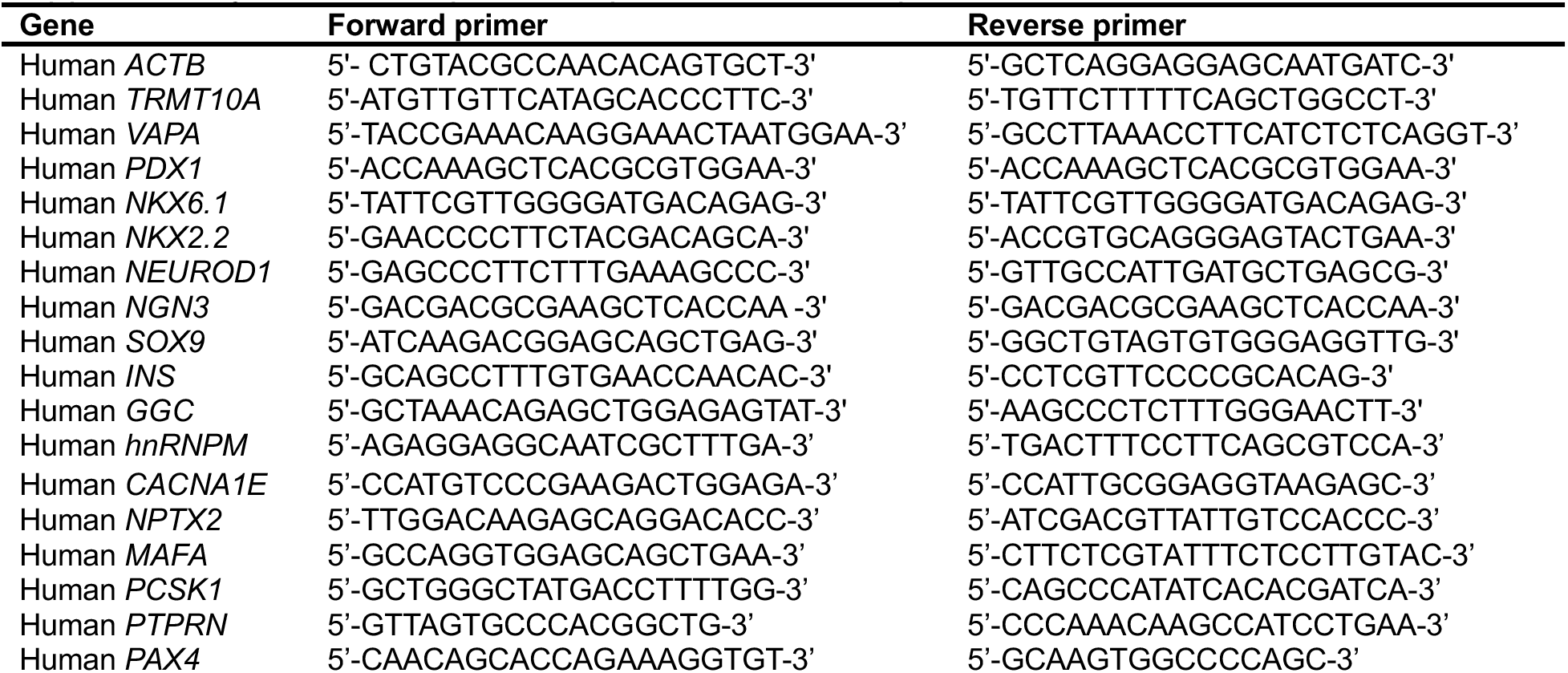
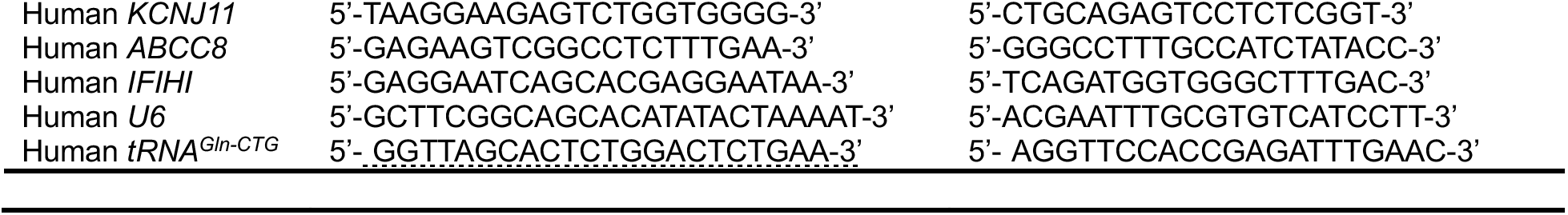
sequence of primers used for qPCR.

**Supplementary Table 10.**

| Composition of complete human islet medium |  |
| --- | --- |
| Component (final concentration) | Brand and catalog number |
| Ham's F10 Nutrient Mix with 1 mM GlutaMAX and 6.1 mM Glucose | Gibco # 41550 |
| 0.75 % BSA Fraction V FFA-Free | Roche, # 10775835001 |
| 1 mM GlutaMAX | Gibco, # 35050-038 |
| 50 µg/ml streptomycin, 50 U/ml penicillin | Lonza, # 17-603 E |
| 50 µM 3-Isobutyl-1-methylxanthine (IBMX) | Sigma-Aldrich, # I5879 |

**Supplementary Table 11:**
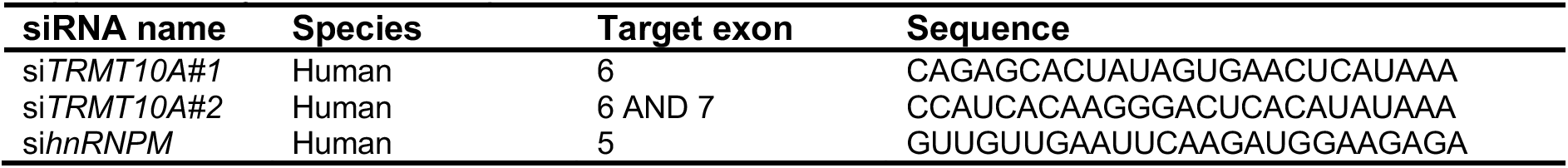
sequence of siRNAs.

**Supplementary Table 12:**
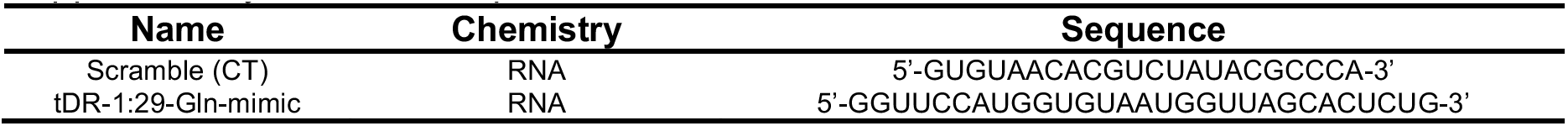
sequence of tDR mimics.

**Supplementary Table 13:**
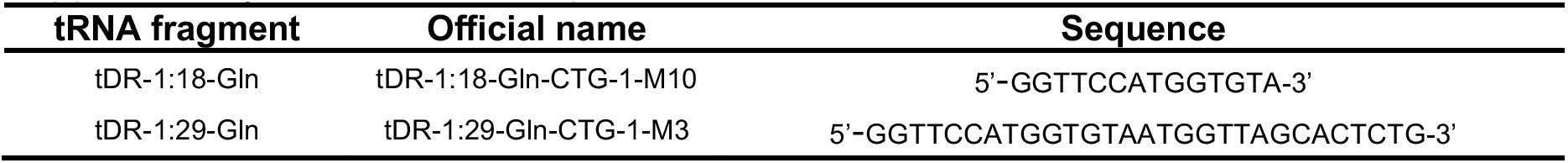
tDR sequence.

**Supplementary Table 14:**
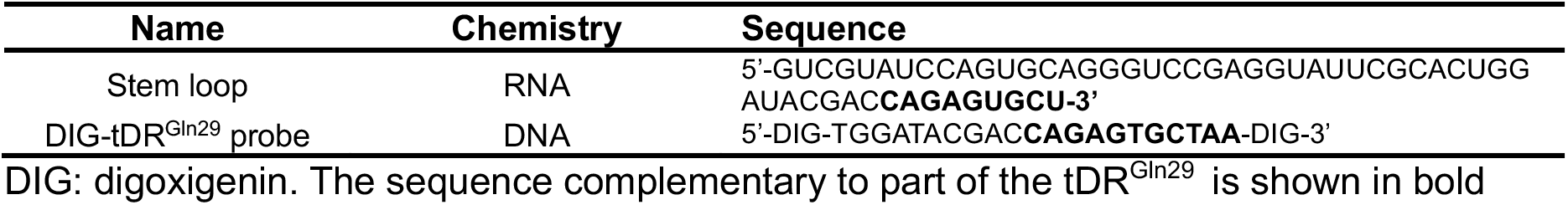
sequence of probes used for LI-FISH.

